# Thalamo-Cortical Interaction for Incremental Binding in Mental Contour-Tracing

**DOI:** 10.1101/2023.12.20.572705

**Authors:** Daniel Schmid, Heiko Neumann

## Abstract

Visual object-based attention marks a key process of mammalian perception. By which mechanisms this process is implemented and how it can be interacted with by means of attentional control is not completely understood yet. Incremental binding is a mechanism required in more demanding scenarios of object-based attention and is likewise experimentally investigated quite well. Attention spreads across a representation of the visual object and labels bound elements by constant up-modulation of neural activity. The speed of incremental binding was found to be dependent on the spatial arrangement of distracting elements in the scene and to be scale invariant giving rise to the growth-cone hypothesis. In this work, we propose a neural dynamical model of incremental binding that provides a mechanistic account for these findings. Through simulations, we investigate the model properties and demonstrate how an attentional spreading mechanism tags neurons that participate in the object binding process. They utilize Gestalt properties and eventually show growth-cone characteristics labeling perceptual items by delayed activity enhancement of neuronal firing rates. We discuss the algorithmic process underlying incremental binding and relate it to the model’s computation. This theoretical investigation encompasses complexity considerations and finds the model to be not only of explanatory value in terms of neurohpysiological evidence, but also to be an efficient implementation of incremental binding striving to establish a normative account. By relating the connectivity motifs of the model to neuroanatomical evidence, we suggest thalamo-cortical interactions to be a likely candidate for the flexible and efficient realization suggested by the model. There, pyramidal cells are proposed to serve as the processors of incremental grouping information. Local bottom-up evidence about stimulus features is integrated via basal dendritic sites. It is combined with an apical signal consisting of contextual grouping information which is gated by attentional task-relevance selection mediated via higher-order thalamic representations.

**Author Summary:** Understanding a visual scene requires us to tell apart visual objects from one another. Object-based attention is the process by which mammals achieve this. Mental processing of object components determines whether they are compatible to the overall object and, thus, should be grouped together to be perceived as a whole or not. For complicated objects, this processing needs to happen serially, determining the compatibility step by step. In this work, we propose a neural model of this process and try to answer the question of how it might be implemented in the brain. We test the model on a case of object-based attention for grouping elongated lines and compare it to the available experimental evidence. We additionally show that the model not only explains this evidence, but it does so also by spending neurons and connections efficiently — a property likewise desirable for brains and machines. Together, these findings suggest which brain areas might be involved in realizing this process and how to reason about the complexity of this computation.

## 1. Introduction

Visual cortex is able to solve a variety of tasks, such as searching a scene for the presence of an object, tracking visual motion, or making visual comparison judgments. While many of these tasks are based on seemingly effortless perception that relies on processes operating in parallel, others require the dedicated deployment of functional resources and only can be executed sequentially [1, 2, 3]. Such selective deployment of functional resources require attentional focus. Endogenous attentional control steers the sequence of visual mental operations. Each sequence forms a dedicated pattern of neural activity and can be used to solve a specific visual task [4, 5, 6]. Object-based attention is one such task, which requires grouping together visual components that may belong to the same object. The goal is to segregate these from distracting parts of a scene, to assemble a visual object, and to disambiguate possible alternatives of explanation [7]. To achieve this goal, two forms of grouping, or binding, can be distinguished: parallel and incremental grouping. *Parallel grouping* automatically extracts matching input patterns into representational elements and groups them by a fixed scheme that determines compatibility based on statistics of co-occurrence. Ambiguities that can’t be resolved by parallel grouping require volitional incremental grouping processes [1]. *Incremental grouping* iteratively and dynamically evaluates compatibility of previously grouped elements with their neighbors. If compatible neighbors exist, they become part of the grouped elements as well and the iterative process likewise continues for their neighbors to discern the assembled shape [8].

Executing an incremental grouping operation serves to bind visual elements via spatial relations not hard-wired as in automatic grouping operations. For example, incremental grouping is deployed to decide upon a spatial relation of two disparate locations in the visual scene. In contour tracing tasks subjects use incremental grouping to determine the end point of a connected line while maintaining fixation of a reference location [9] (Fig. 1). Execution of contour tracing requires four phases of processing [12]. During the first phase, features are detected by an onset response. The second phase establishes a *base representation* of foreground and background elements by parallel grouping. In contour tracing scenarios this base representation encodes bottom-up evidence for the existence of line elements that match the respective neuron’s retinotopic position and feature selectivity. During the third phase, an *incremental representation* is computed by combining the existing bottom-up evidence with an attentional binding signal into elements of an object. After the process concluded, during a fourth phase a decision is made based on the integrated evidence about the visual objects. The formation of the incremental representation during the third phase relies on incremental grouping to propagate the binding signal and iteratively group the attended object [8]. The binding signal consists of two kinds of information: contextual information and task-related information. *Contextual information* stems from a neuron’s neighborhood and serves to evaluate *compatibility* between the features a neuron and its neighbors encode. *Task-related information* signals whether this compatibility is behaviorally relevant and provides the attentional seed point for the incremental representation. This seed is selected, or indexed, by the fixation point in contour tracing panels.

**Figure 1:**
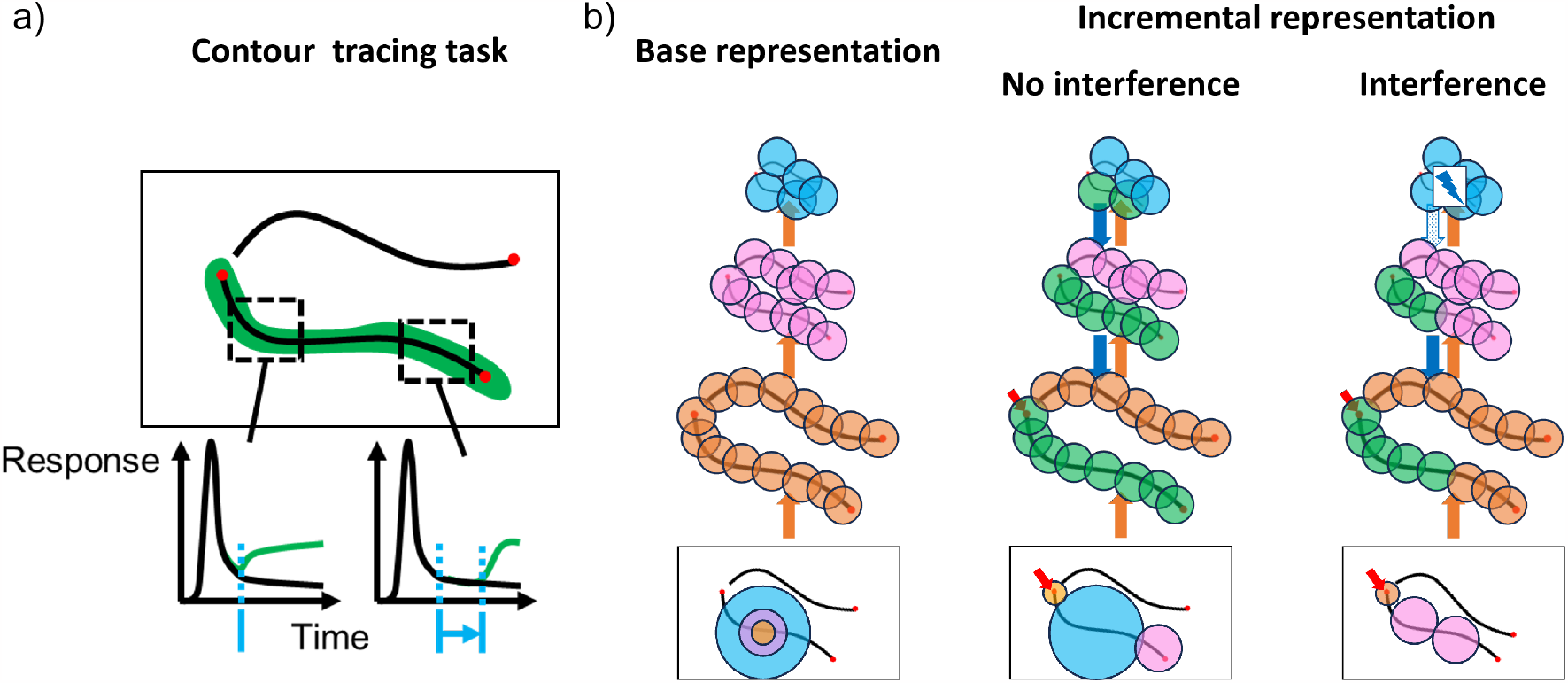
Incremental binding in contour tracing tasks and the growth cone hypothesis. **a**) During contour tracing tasks a fixation point is presented together with a pair of contours. Subjects need to determine which contour is connected with the fixation point and perform a saccade towards the contour’s end point in response. Neurons coding for the connected contour show an up-modulation of their activity (green) with respect to a baseline response profile (black) [9]. The onset latency of this up-modulation is thereby dependent on the distance among the line from the fixation point (blue). Furthermore, the up-modulation is found to be stationary over time, so that ultimately the whole contour representation is up-modulated. These findings indicate a serial processing of incremental binding after an initial processing phase of the stimulus. **b)** A specific hypothesis of incremental binding is the growth-cone hypothesis [10, 11]. It explains how speed-variations in incremental binding arise from the distance to distracting elements in the vicinity of the target object. The initial processing phase has extracted features of the stimulus in parallel to build a hierarchical base representation (left). Within this hierarchy incremental binding starts from an attentional seed position (red arrow) and can proceed at different scales. While coarser scales span more space and can proceed faster (“no interference” case, center), they might suffer from interference with close-by elements. In the case of interference finer scales still can perform incremental binding, but with a reduced speed (“interference” case, right). Neural receptive fields are represented by colored circles as an overlay to the input image and are drawn with their relative sizes. Colors are matching the respective level of the hierarchy on which the neurons are localized. Green colored circles denote neurons that are up-modulated and have become part of the incremental representation. For the same amount of time, more neurons can be grouped in case higher, coarser scales can participate in incremental binding. The red arrow in the input displays denotes the location of the seed of attention, which triggers the execution of the execution of the incremental binding process.

In order to build such incremental representations different neural coding schemes have been proposed [7]. To make the presence of an object explicit, one candidate scheme instantiates all possible combinations through hard-coded feature selections.

Such coding requires an expensive realization in terms of the number of neurons and their connections. Alternatively, binding through synchronization of neural oscillations and the delayed up-modulation of neural firing rates have been proposed as other schemes [13]. The latter frameworks are more flexible and more efficient to implement, particularly utilizing those neurons which are already involved in the base representation. These can be dynamically recruited for the incremental assignment of task-related context linking into perceptual objects. The evidence from neural correlates of the incremental binding mechanism indicates a constant up-modulation of neural firing rates across the whole attended object [9, 7]. This further discerns object-based attention by incremental binding from the idea of an attentional spotlight, where only one proportion of the object would be up-modulated at a time [14]. Additionally, the onset time of the up-modulation depends on the neuron’s distance along the curve [9]. This is taken as evidence that an active labeling process operates to incrementally propagate the binding signal along the object.

Does such contour labeling operate independent of the context of the scene? Further investigations showed that the speed of incremental binding is dependent on the target object’s distance to distracting neighboring elements and that this behavior is best explained in terms of the so-called *growth-cone hypothesis* [10, 11]. The growth-cone hypothesis incorporates the observation that incremental binding is a scale-invariant process [15, 10, 11]. The local speed of incremental binding is determined by the neuron with the largest receptive field, that doesn’t interfere with distracting elements in the scene. This way, incremental binding speed is high if the relative distance to distractors is large and it slows down if close-by distractors interfere for neurons with larger receptive fields leaving binding signal propagation to neurons with smaller receptive fields (Fig. 1).

Incremental binding has been investigated on all levels of analysis [16]. On an *implementational level* models were proposed, of which some are mechanistic and some are rather phenomenological. The different models address different neuroanatomical scales and range from a laminar circuit level [17], over the level of local pools of neurons representing the computation by predefined interactions [18, 19, 20], to the level of neural networks exhibiting learned interactions to perform incremental binding tasks [21, 22] or to match behavioral evidence [23, 24]. On an *algorithmic level* more abstract descriptions either discuss incremental binding directly [4, 25, 26, 8] or its embedding in cognitive models [6]. Likewise the *computational level* of incremental binding is discussed in terms of visual routines and cognitive programs [4, 5, 6] and provides connections to theories about binding in general that embed the computation within the broader context of object-based visual processing [27, 28, 29, 30, 31, 32, 33, 7]. Yet, integrative models that link across the levels of analysis are scarce. This current state of findings motivates the following questions: Are there generic principles for mechanisms of incremental binding that also help to better analyze and compare existing models? Is there a normative account that can guide a choice of minimal elements to implement such mechanism? And, what predictions about binding in the brain would arise from a model which adheres to such generic and normative principles?

In this work, we present a mechanistic model for incremental binding that spans multiple levels of explanation and links them to a normative account of connection and representational complexity. Thus, it spans descriptional levels from local pyramidal cell computations to inter-areal interactions and utilizes only a minimalistic set of computational elements and connectivity patterns. Through extensive simulations, we show that the model explains a broad range of experimental evidence. Furthermore, we analyze the model considering functional and complexity constraints in a framework of incremental binding. In this more functional investigation, we consider several architectural principles of distributed neural processing that could implement the desired computational operations for incremental binding. Furthermore, the serial composition of multiple elemental operations in a mental routine defines additional constraints. We argue that the wiring and, thus, representational network complexity, can be utilized to assess the efficacy of a specific network structure and its intersection of components. We show the network architecture proposed in this contribution possesses favorable properties regarding the complexity of its connectivity and neural code. Considering the simulation results, the model architecture, and recent experimental evidence about functional anatomy, we then suggest coupled cortico-cortical and cortico-thalamo-cortical loops as a likely target circuitry that could implement such incremental binding mechanism in the brain.

## 2. Results

Based on the motivations above, we now present a mechanistic model architecture that implements neural incremental binding operations. Through numerical simulations, it explains a broad range of experimental evidence using only a single set of model parameters. An additional theoretical consideration relates the model to other possible solutions of incremental binding and reveals its favorable properties regarding model complexity.

### 2.1. The incremental binding model

The resulting model implements experimental findings about incremental binding and offers a mechanistic explanation by means of a dynamical neural network architecture (Fig. 2). The model is described at a mesoscopic level, where neural processors resemble the computation of a local neural population. The neural processors capture functional principles of pyramidal cell computation and utilize a rate-based encoding scheme. The architecture is organized by modules. It consists of a visual cortex module, an interfacing module and a task module. The *visual cortex module* is organized in a hierarchy of retinotopic feature maps, which represent oriented contrast information in a scale-space pyramid. Feature maps order neurons in two-dimensional sheets. There, neurons are placed at retinotopic locations in a rectangular grid, code for specific stimulus features and exhibit local connectivity patterns. The visual cortex module computes the base representation as well as the incremental representation and propagates information among feed-forward and feedback connections from lower layers to higher layers and vice versa. The *interfacing module* is likewise composed of neurons in a two-dimensional sheet. Yet, different from the visual cortex module, the neurons don’t encode for multiple stimulus features, but only for whether a retinotopic position is part of the attended object or not. The interfacing module serves to interface task-related information between the task module and the different hierarchical layers of the visual cortex module, and broadcasts information about the binding state originating from one of the hierarchical layers to all other ones. Due to its connectivity it forms a shallow, rather than a hierarchical interaction with the visual cortex module and is, thus, associated with higher-order thalamic regions [34]. The *task module* represents an attentional seed location. By this seed location the task module performs the selection of from where the incremental binding operation should start. This selection happens by forwarding the location to the interfacing module. There, it constitutes the first entry in the sheet and, thus, determines the starting point from where an object should be incrementally grouped by attention. Such selection process is necessary to decide which object should be grouped and can likewise be useful in disambiguating the representation.

**Figure 2:**
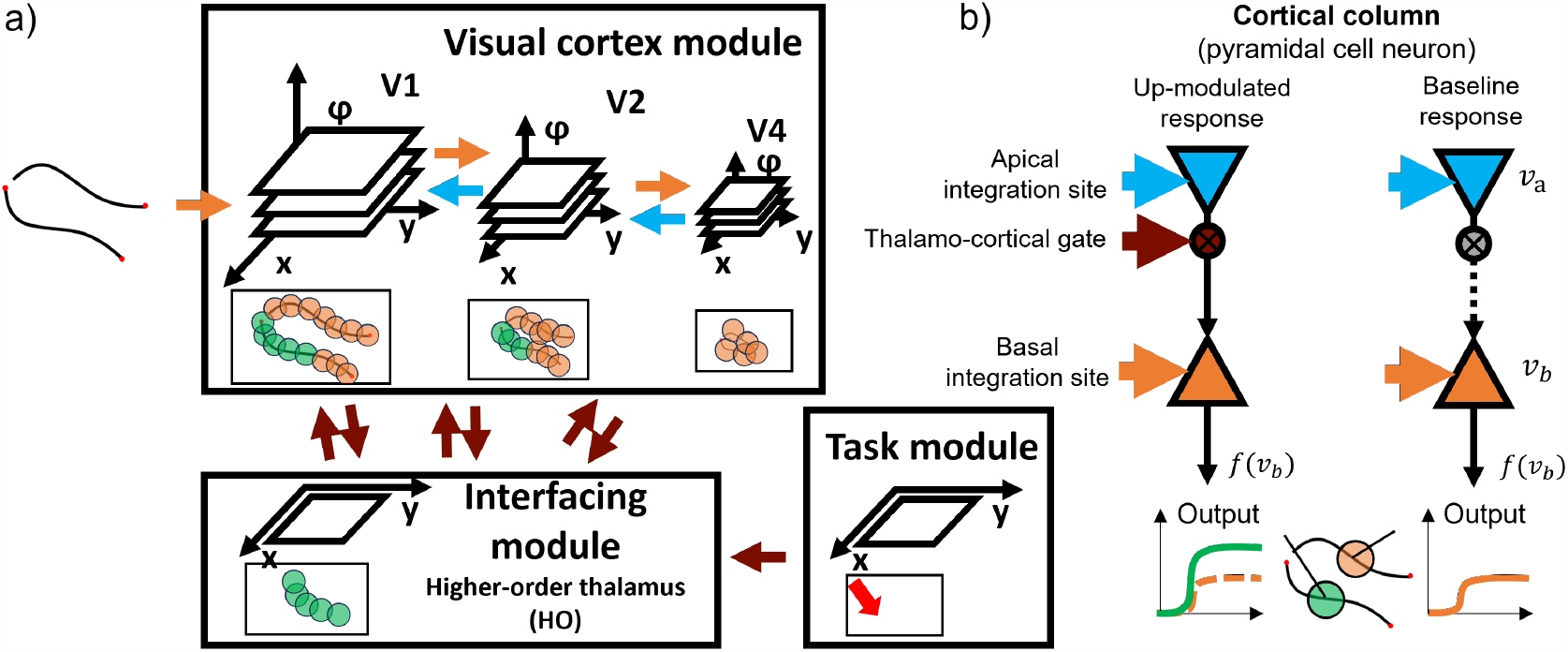
Thalamo-cortical model architecture for incremental binding with local grouping processors that are represented by cortical columnar pyramidal cell computation. **a**) The proposed incremental binding architecture consists of three modules. The visual cortex module encodes the stimulus in a high-dimensional feature space organized along a spatial hierarchy. Adjacent hierarchical layers interact recurrently via feedforward (orange) and feedback (blue) connections. The interfacing module encodes task-relevant spatial locations. It is reciprocally connected with the visual cortex module aggregating information per location across features and projecting information back onto a local neighborhood irrespective of feature expression and hierarchical scale. The task module provides external input to the interfacing module to set the initial seed point of attention. It is proposed that these modules have biological counterparts in visual cortex areas V1 to V4 (visual cortex module), higher-order visual thalamus (interfacing module) and regions involved in attentional selection in either prefrontal, parietal, or temporal cortical areas (task module; the red arrow denotes the location for attentional selection). **b)** Neural computation of cortical columns within the visual cortex module is represented by dynamical, rate-based pyramidal cell models consisting of two compartments and three types of input. From the coincidence of these inputs local grouping information is evaluated to change the neural activity from a base representation state into an up-modulated activity state coding for binding information. There, feedforward signals about the base representation arrive at the basal compartment and can drive the cell into a baseline level activity (orange), while feedback signals arrive apically and are combined with thalamo-cortical gating signals before taking an up-modulating effect on the basal computation. This up-modulated state of activity represents the binding state (green).

Within the visual cortex module, each retinotopic position is represented by a *pyramidal cell model neuron*. These model neurons abstractly capture the computation of a cortical column. Each neuron engages in the computation of the base representation and the incremental representation. Model neurons possess to sites of integration, a basal dendritic site and an apical site. At the *basal site* input-related information enters via feed-forward signal propagation and is required to compute the base representation. This base representation results in a neuronal activity level that encodes the likelihood of feature presence. At the *apical site* contextual information via feedback signal propagation from higher layers and binding signal information from the interfacing module enter. In case both kinds of information coincide, the apical site relays this signal to the basal site. There, it can be integrated with the base representation to compute the incremental representation. Thus, the incremental representation depends on the co-incidence of basal and apical signals of a model neuron. As a result, the neuron’s activity level will be up-modulated and shifted towards a regime of higher activity. Such up-modulated activity distributions represent the labeling of neural elements to signal binding through enhanced firing rates. The interfacing module aggregates firing rate activity from among the pyramidal cell neurons of the visual cortex module. It filters these rates for such higher activity level to represent the binding state at that retinotopic position. The label is then broadcast back to spatial neighbors across scales in the visual cortex module to further propagate the binding signal locally.

### 2.2. Simulation results capture experimental evidence about incremental binding

#### 2.2.1. Tracing time depends on contour length and captures neural correlates

To investigate the incremental binding capabilities of the model we evaluated it on contour tracing tasks similar to those experiments conducted with human and nonhuman primate subjects (Fig. 3). The model is able to capture the main findings and neural correlates. The model neurons encode the binding state by a constant up-modulation of their firing rate. The incremental nature of this binding process becomes visible from the time course of this up-modulation. After an initial onset phase, neurons located along the attended curve become up-modulated. There, the modulation onset time depends on the neuron’s distance along the contour from the seed point of attention. Once all neurons along the contour have been bound, the representation reaches a stable equilibrium and the incremental binding mechanism comes to rest. These findings likewise imply, that the time to incrementally bind a curve depends on the curve length. Therefore, decisions about the connectedness of two points will take longer the further apart these points are along the curve. This is in line with experimental findings not only on a behavioral level [35], but also regarding the neural correlates of incremental binding [9].

**Figure 3:**
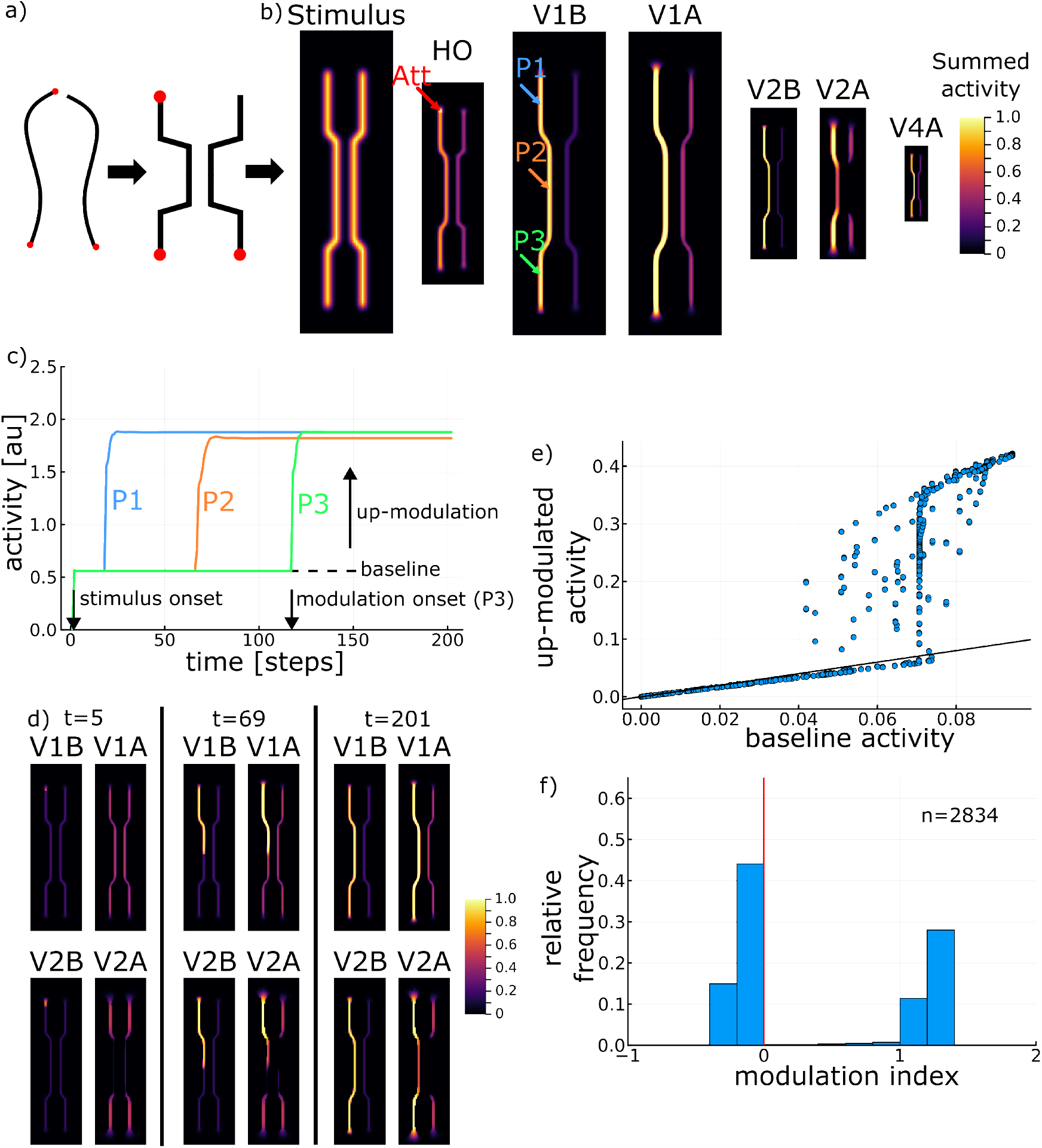
Simulation results for stimulus with varying target-distractor distance. **a**) The synthesized stimulus captures the main properties of contour tracing stimuli [10] by providing a varying distance between target and distractor contour. **b)** Neural representation after incremental binding has taken place for basal (VxB) and apical (VxA) activity across layers V1 to V4 and higher-order thalamus (HO). Arrows denote the attentional seed location (Att) and probe locations (P1 to P3). Activity for layers V1 to V4 is summed across the feature channel and normalized to 1.0. Layer sizes reflect their scale space ratios. **c)** Neural activity across time at the probe locations shows the main hallmarks of incremental binding, i.e., a constant up-modulation when being bound and a distance-dependent modulation-onset latency. **d)** Neural representations at the beginning of (left), during (center), and after (right) incremental binding. While apical context of V1 is available for the complete contour from the beginning, the context of V2 is missing at first for the narrow contour segment due to interference. **e)** Distribution of neural activities along the contour while being grouped in comparison to their baseline activity. **f)** Distribution of modulation indices for neurons along the target contour.

#### 2.2.2. Tracing operates on multiple scales - as predicted by the growthcone hypothesis

Consider once again two curves among which one needs to be selected and incrementally bound via attentional selection. As detailed above, the activity in the underlying neural contour representation is up-modulated to enhance the neurons’ firing rate [7]. It was demonstrated [35] that the time to mentally trace a contour depends on the distance between the target and the distractor curve. The growth-cone model of object-based attention predicts that neural spreading can progress at larger scales when contours are positioned farther apart [10, 11]. The hypothesis is that the faster spreading is accomplished by cells having larger receptive fields operating on a spatial grid that is sampled less densely by the neurons that perform the spreading. The spreading mechanism then selects the largest scale on which neurons represent the target contour while not interfering with other distracting contours captured by the same receptive field. For densely spaced contours, only cells with smaller receptive fields do not interfere and are able to resolve the spreading unambiguously [36]. The growth-cone behavior realizes a scale-invariant incremental binding process [10, 11], a property of incremental binding that was as well behaviorally identified by earlier experiments [15].

To investigate this function in the proposed model, we more specifically performed experiments with respect to the growth-cone hypothesis of incremental binding. Thus, to vary the degree of target-distractor interference along the contour, we systematically varied the proportion of the overall stimulus which has a more narrow distance between the target and distractor curve (Fig. 4). In line with the idea of multi-scale incremental binding, the time until the whole curve is bound together depends on this narrow-wide proportion. The longer the narrow section of the configuration the more time it takes the model to reach the other end of the curve and vice versa. This tracing time result already indicates that the speed of the model’s incremental binding computation depends on the target-distractor distance.

**Figure 4:**
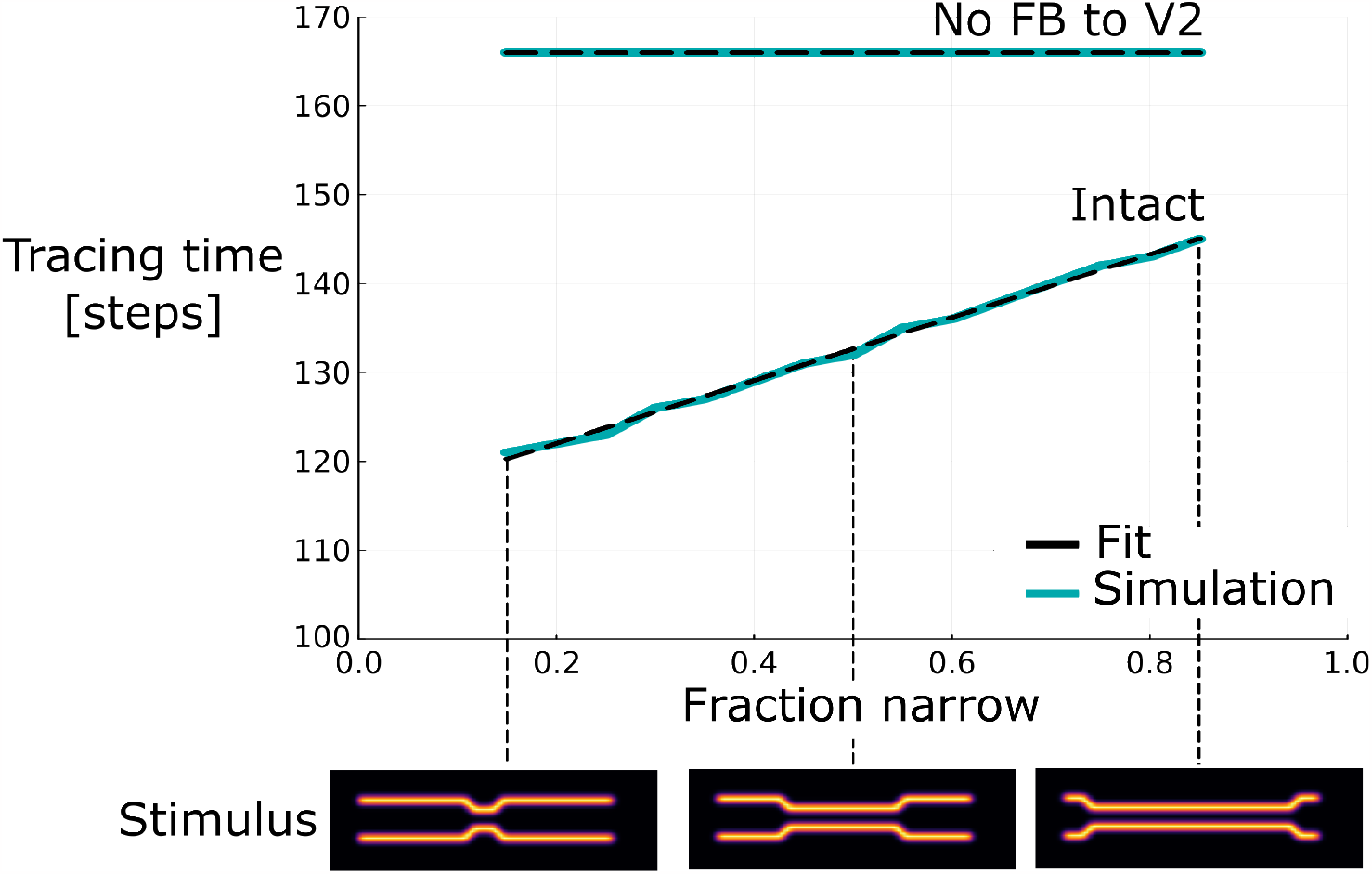
Tracing time across input stimulus conditions of different distance proportions between target and distractor curves. Tracing times for different proportions of narrow distances among target and distractor line segments for the intact model and a model without feedback connections from V4 to V2 (top) and representative stimuli for different such ratios (bottom). The tracing time dependency on the fraction of narrow distances along the contour is well explained by a linear model (black, linear fit; blue, simulation data). In case of lesioning feedback connections of the higher hierachical scales of the model no such distance-dependent tracing time variation is observed anymore. Instead, the tracing time becomes constant. Results extend a previous investigation that was performed with a simpler model version [20].

In addition to measuring the overall time to complete the binding process, the time it takes to label a segment of the contour can be computed. If the length of the respective contour segment is known in addition, then this yields an estimate of the binding speed for each segment separately. From these estimates a distance-dependent variability of binding speed can be seen from the modulation onset times of model neurons along the contour (Fig. 5). There, a reduction of binding speed coincides with the narrow section of the curve configuration (2.14 neurons/step, 1.63 neurons/step, 2.12 neurons/step, for the first wide segment, the narrow segment and the second wide segment, respectively). This observation is likewise reflected by the apical integration process along the hierarchical scales (Fig. 3 d). While apical context information on the higher hierarchical scale can be established for the wide contour segments right from the beginning, this context information is only becoming available as the binding progresses for the narrow segments. Conversely, on the lower hierarchical scale this apical context information is established also for the narrow segments right away and readily can be integrated with the basal signal once a coinciding binding signal from the interfacing module become available. Thus, it can be seen in the model how this distance-dependent speed variation links to the apical representation of contextual feedback information.

**Figure 5:**
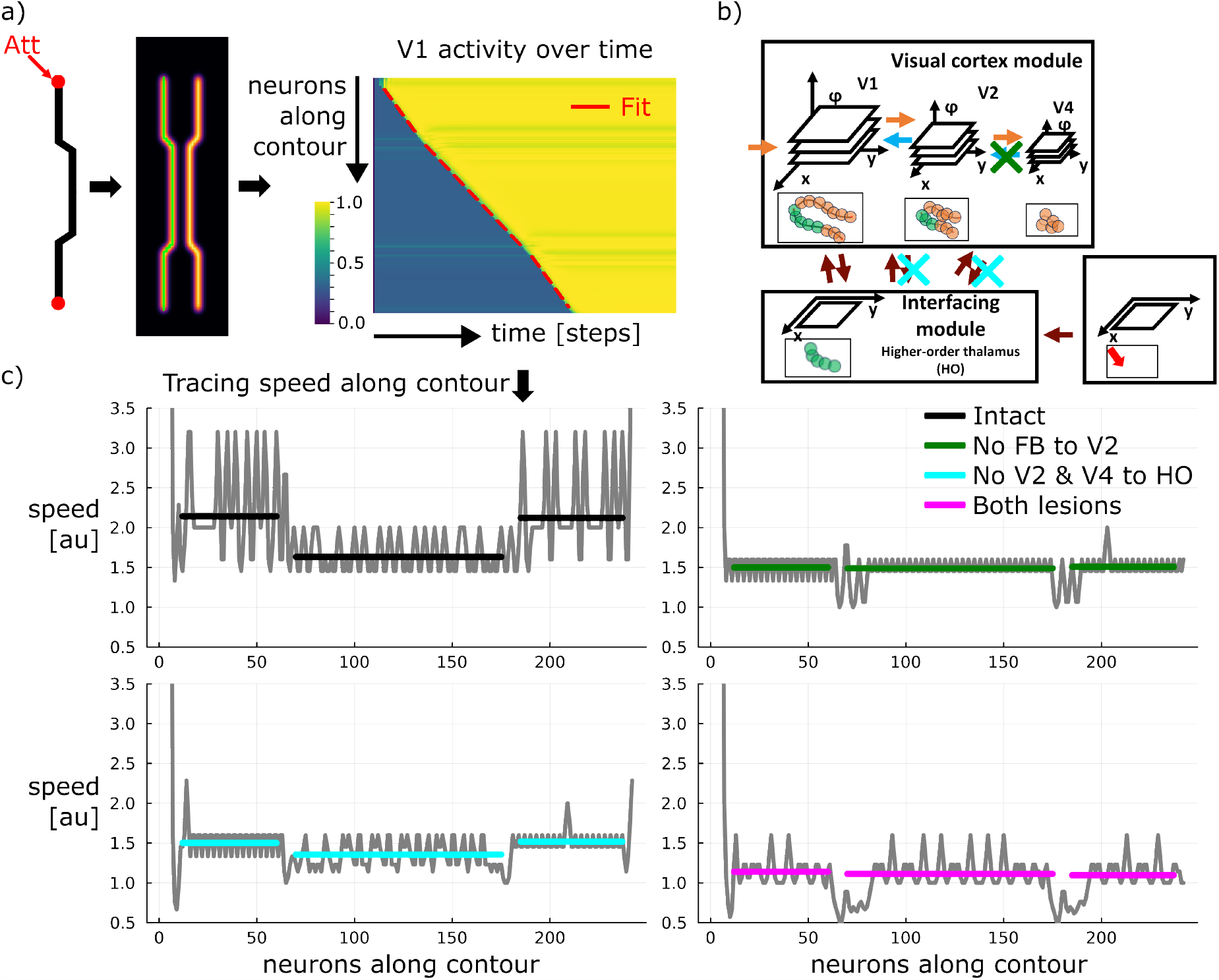
Tracing speed varies along contour dependent on the lateral distance to distractor elements. **a**) Neurons along the contour show an modulation onset latency dependent on their distance from the attentional seed location. For the contour segments of constant distance to the distractor linear regressions provide an excellent fit to the onset latency along the line. The green line overlaid on the stimulus denotes the path along which neurons have been read out. V1 activity has been summed across feature channels and normalized to the maximum per neuron. **b)** Based on the modular architecture specific connectivity patterns can be lesioned to investigate their functional role. **c)** Computing smoothed finite differences between neighboring onset latencies provides an estimate of the tracing speed profile along the contour. Transforming the regressed latency slope into an average speed per segment provides a good fit to the speed profile. Lesion studies yield speed profiles differing from the intact model version. The differences distinctly depend on the type of lesion. The different lesions are indicated in the architectural diagram. Gray lines denote speed estimates, black denotes the fit for the intact case, colored lines denote the respective fits for the different lesion cases.

#### 2.2.3. Ablation studies showcase model predictions

The results so far established that the model is able to implement incremental binding and that these binding capabilities exhibit growth-cone-like variations in binding speed. There, the binding speed variations coincide with a variation in the apical contextual state of neurons along the hierarchy of the visual cortex module. What cannot be seen directly from the results so far, is whether there is a causal link between the introduced scale space by the hierarchy and the binding speed variations, or whether the variation in binding speed arises within the model dynamics independently of the different scales. For that reason, we performed ablation studies to learn more about the functional relevance of the model’s components. In order to selectively impair the complex information flow between the model’s components, we eliminate connections. We investigated the effect of removing either specific feedback connections within the visual cortex module or specific connections between the interfacing module and the visual cortex module (Fig. 5).

For investigating feedback connections we removed the connections on the larger scale, i.e., from the highest layer (V4) to the middle layer (V2). Under this ablation condition the speed profile becomes near-constant along the curve’s outline and, thus, misses the drop in speed for the contour segment with a narrow target-distractor distance (1.50 neurons/step, 1.49 neurons/step, 1.51 neurons/step). Furthermore, the constant speed level in this ablation case resembles rather the speed of the narrow than the wide segment of the contour’s outline from the intact case (1.63 neurons/step vs. about 2.13 neurons/step, cf. above). Likewise, this constancy in binding speed can also be seen at the level of the total binding times, which don’t vary substantially with the proportion of narrow segment length (Fig. 4).

This lesion cases indicates that the model’s feedback are necessary to exhibit target-distractor dependent speed-variations. Thus the model’s growth-cone behavior can’t exist without the coarser scales’ feedback interaction. Also, it can be seen that the target-distractor interference for the narrow segment during the intact case has nearly the same effect as removing the higher-layer feedback during the lesion case, since both of these causes result a similar speed. That the interference for narrow segments leads to feedback that is virtually absent at the higher layers can also be seen again from the initial apical representation of the intact case (Fig. 3 d). This observation is further evidence for that the model implements growth-cone behavior by feedback among spatial scales and the interference thereof in case of close-by distractors.

For investigating connections between the interfacing module and the higher layers of the visual cortex module we removed connections from the middle layer (V2) and upper layer (V4) to the interfacing module. While an overall drop in binding speed can be observed, the qualitative speed profile stays to a degree intact exhibiting higher speeds for the wider segments of the contour’s outline and a further reduced speed for the narrow segment (1.50 neurons/step, 1.36 neurons/step, 1.52 neurons/step).

The interfacing module accumulates evidence about up-modulated activity from all the layers of the visual cortex module and gates the subsequent binding increments. Thus, removing some of the inputs to this evidence accumulation, i.e. inputs from V2 and V4, slows down the process. As the inputs of larger scales affect a bigger neighborhood than inputs from V1, not only the overall process is slowed down, but also part of the relative difference in speed between wide and narrow segments is lost. So, the only scale on which evidence can be accumulated is the scale of V1. Yet, once enough evidence is accumulated from V1, a speed-variation is still noticeable, since the contextual integration of feedback from V4 to V2 is still intact. Once the interfacing module opens the gates and sends a task-relevance signal to the visual cortex module it allows also this higher-level (V4 to V2) integration to take place, which, in turn, affects the lower-level (V2 to V1) integration.

Applying both lesion cases together, i.e. removing feedback connections from V4 to V2 and connections from V4 and V2 to the interfacing module, results in a constant speed profile again. Notably, this constant speed is lower than any of the speeds for the individual lesion cases (1.14 neurons/step, 1.11 neurons/step, 1.10 neurons/step). This lesion results in effectively decoupling the higher layers, and thus, scales, from taking part in the incremental binding process: higher-level contextual integration cannot occur due to missing feedback from V4 to V2, and neural activity in V2 and V4 cannot affect the evidence accumulation in the interfacing module due to missing connections. Thus, only V1 will be able to integrate contextual evidence from V2 and enter it into the interfacing module. Consequently, this lesion realizes the case of incremental binding on a single scale.

Together, we interpret these results as follows. The lesions indicate that both kinds of connections influence the dynamics and speed of binding. Yet, specifically the feedback connections are responsible for the multi-scale effects associated with growthcone-like incremental binding properties. Conversely, projections to the interfacing module affect the integration process overall and lead to a reduction in overall speed, if the projections are missing. In sum, a causal relationship exists between the model’s feedback connections in scale space and it’s growth-cone-like binding properties.

#### 2.2.4. Task-dependent Gestalt disambiguation is based on attentional seeding

So far the model has been tested on configurations with contours adjacent to each other. These configurations already revealed the model’s incremental binding properties and it’s growth-cone-like behavior. There, the target contour was selected by the seed of attention on one of its ends which served as the fixation point. The influence of the other, distractor contour on the target contour’s incremental binding process was mediated by the distance between the contours. What remains open from these investigations, is what happens to the model’s binding process when target and distractor object intersect. Therefore, we chose a configuration with two contours crossing each other, similar to the stimulus used in [9].

We probe the model presenting pairs of contours which cross each other at about half the length of each contour (Fig. 6). The attentional label again grows along the target contour until it reaches the location of the crossing point. Geometrically, the attentional label has three options to spread along one of the continuing arms that leave the junction or even tag multiple contour segments by splitting the attentional signal. The simulation demonstrates that the attentional label spreads along the smooth continuation of the contour at the entrance of the crossing. In other words, the incremental binding follows the Gestalt law of good continuation [37]. These grouped contours match the target objects from [9]. We emphasize that mechanisms of incremental grouping in the model recruit mechanisms that are also used for linking perceptual items in base grouping operations.

**Figure 6:**
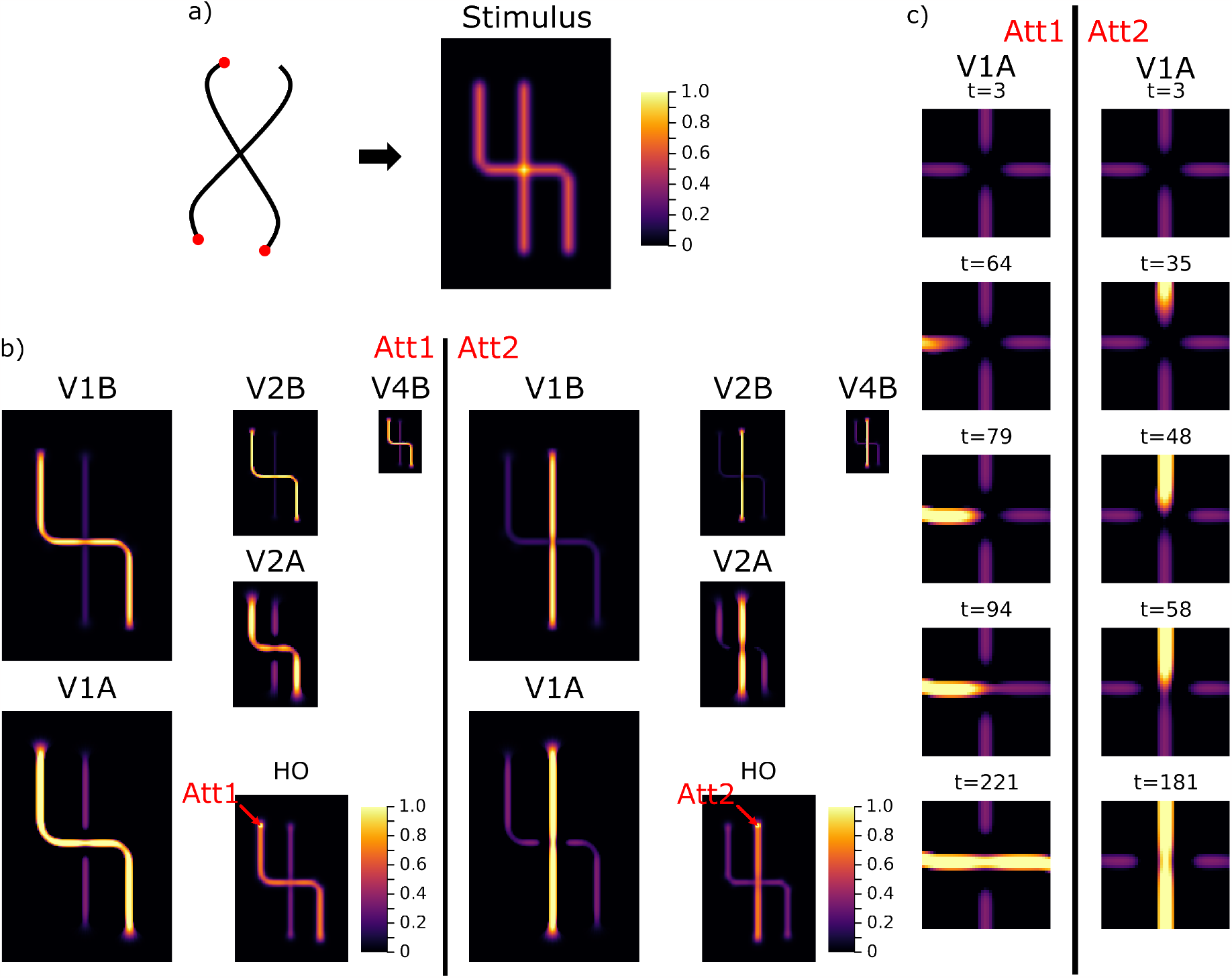
Task-dependent grouping of an intersection. **a**) The synthesized stimulus captures the main properties of intersection stimuli [9]. The fixation point location can be varied between the two contours and steers the seed of attention to indicate which of the contours should be grouped into an object (Att1, Att2). **b)** Dependent on the attentional seed location the simulation results in either one or the other line being grouped. The behavior follows the Gestalt laws of good continuation. **c)** Over the time course of integration the role of the apical representation becomes visible. After the base representation has been formed the intersection is initially not represented apically in either condition (Att1 and Att2). This indicates no compatibility to either contour segments, because of mutual inhibition between the feedback projection to orthogonally oriented feature maps. Only once the up-modulation signal reaches the vicinity of the intersection, the mutual inhibition becomes biased and allows the intersection elements to become compatible to the respectively grouped line elements.

By investigating the model’s internal representation one can learn about why the binding happens like this (Fig. 6). After the initial base representation has been formed by the model’s feedforward processing, the apical representation around the crossing point exhibits a configuration with no preference for compatibility to any of the curve segments. Within the representation of the visual cortex module, neurons project to neurons in the next lower layer. There, they target neurons that are selective to similarly orientated contrasts via excitatory connections, and neurons that are selective to orthogonally oriented contrasts via inhibitory connections. The evidence for the four segments surrounding the crossing are equally strong encoded in the base representation. Thus, neurons at the location of the crossing, that are selective to either horizontally or vertically oriented contrasts receive the same amount of excitatory and inhibitory feedback projections from the likewise and orthogonally tuned neurons of the upper layer, respectively. As a consequence, the initial apical representation at the crossing location is balanced out for horizontal and vertical preference. These equal contributions only change later on, once the incremental binding signal spreads into the vicinity of the crossing point’s representation. Then, neurons coding for the orientation of already bound segments are up-modulated. Their enhanced activity is subsequently fed back onto their lower layer neighbors and biases their apical input towards oriented contrasts collinear to those of the already bound curve segments. Through this interaction, the model establishes an incremental binding version conforming not only with neural correlates of incremental binding and the growth-cone hypothesis, but also with the Gestalt law of good continuation in the crossing contours case.

### 2.3. Complexity Considerations of Incremental Binding Architectures

The proposed model on an implementational level links descriptions from pyramidal neurons to inter-areal computation and provides a dynamical realization of the incremental binding algorithm. Beyond the simulation results that show that the model is able to explain a set of experimental evidence, a theoretical analysis of the model’s complexity yields insight into how far it fulfills a normative account. To this end, we analyze the model’s complexity in terms of connectivity and representation and compare it against alternative realization schemes for solving incremental binding. This analysis extends on the canonical framework of incremental grouping by Roelfsema [8] building upon earlier investigations of our own [21, 20].

#### 2.3.1. Basic components for incremental binding

The analysis focuses on incremental binding models that capture the neurophysiological evidence discussed above. Thus, models need to be able to compute a base representation and an incremental representation, and they need to do so within a hierarchy of multiple scales to achieve growth-cone-like incremental binding. Abstractly, such models can be described by network graphs [8]. These network graphs are composed of two basic elements: neural processors (NProcs) and channels. *NProcs* capture the computation that takes place within a local microcircuit at a certain retinotopic position and hierarchical scale. *Channels* capture the network’s connectivity structure and form passive elements that relay information between NProcs unidirectionally. Different models of incremental binding can be described by these elements. Such models can be distinguished by the properties of the network graph, i.e., by how NProcs encode the representational information, what types of channels exist, and how they connect NProcs with each other.

Such distinction between models helps categorizing them with respect to their computational properties and complexity. A comparison based on the complexity properties additionally answers the question of to what extent a model not only provides a mechanistic account for incremental binding, but also normative one.

The following subsections investigate the complexity of individual network components in more detail. First, different variants of how NProcs can encode the necessary information are discussed with respect to their representational complexity. Second, it is investigated how different channel types can be used to provide NProcs with the necessary information for their computation and what it means in terms of connectivity complexity. Third, it is shown which possibilities exist to interface with these networks in terms of task signaling and read-out and what are their pros and cons. The last subsection then interrelates these insights to the specific model proposed in this work.

#### 2.3.2. Local neural processors and their representational complexity

To perform incremental binding, models need to adhere to the main phases of the incremental binding algorithm, i.e., computing a hierarchical base representation at first and an incremental representation afterwards. Each NProc is involved in this computation locally and needs to implement according decision rules. A symbolic interpretation of these decision rules provides an understanding of the required flow of signals within the network. On a local NProc level this computational logic can best be understood in terms of boolean logic using “and”(*∧*) operations. Similarly, a population level view across NProcs can be used to describe the incremental binding algorithm. There, set-theoretic terms describe the algorithm, where becoming a part of the set of NProcs coding for a specific variable would translate into a union (*U*) operation. In this contribution the necessary computational logic will be described in local boolean terms to understand the decision functions each NProc needs to implement locally.

For computing the hierarchical base representation NProcs need to have access to the base evidence provided by the respective lower hierarchical level (or by the visual input directly in case of the lowest level’s NProcs). If enough evidence is provided to the respective NProc it should likewise code for this evidence at its output. In boolean terms, each NProc that is part of the base representation computes a binary variable about the evidence for base representation at its position *X*_*Base*_ *∈ {noevidence, evidence}* as a function of the incoming base representation variables 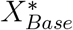 from other NProcs 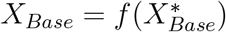.

To compute whether an NProc belongs to the incremental representation or not it needs to have access to two distinct signals: the base representation for the space-feature combination for which it is selective at the respective hierarchical scale, and the incremental representation from its neighboring NProcs. If enough evidence for the local NProc’s base representation is available and the incremental representation among its neighbors provides a compatible constellation for this NProc, then it should likewise become a part of the incremental representation and code for this circumstance at its output. In boolean terms each NProc that is part of the incremental representation needs to compute a binary variable about its binding state, i.e., *X*_*Binding*_ = *{unbound, bound}*. This variable is computed by a logical “and” between the NProc’s base evidence *X*_*Base*_ and the evidence about the incremental representation, which is a function of the binding state variables 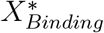 of the neighboring NProcs 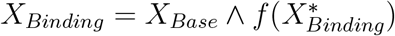.

Different alternatives exist to implement this boolean NProc logic: a naive *two-node static* implementation and a *single-node static* implementation which employ a static encoding, and a *single-node phasic* implementation which relies on bi-phasic codes.

A naive *two-node static* implementation of these decision rules above will require *log*_2_ 4 *bit* = 2 *bit* to encode the two binary variables *X*_*Base*_ and *X*_*Binding*_ as independent output signals. In this case a model either assumes to have two NProcs per space-feature combination with a binary output code for 2 *×* 2 distinct values, e.g., one coding for the base representation and one coding for the incremental representation, or to have one NProc that provides a more complex output code with 4 distinct values to cover all combinations between the variables. An improved *single-node static* version considers the fact, that the binding state *X*_*Binding*_ is dependent on the base representation *X*_*Base*_, in such a way that it can never be in binding state if no base evidence exists for that space-feature combination, i.e., *X*_*Binding*_ = *bound* if *X*_*Base*_ = *noevidence* is an impossible state. Thus, effectively only 3 distinct values *{noevidence, evidence ∧unbound, evidence ∧bound}* need to be encoded at the output. As the binding states implies the existence of base representation evidence the value can simply become *{noevidence, evidence, bound}* (Fig. 7 a). This leads then to *log*_2_ 3 *bit ≈* 1.58 *bit*, that could be encoded by a single NProc with a ternary output code. Another alternative exists in the *single-node phasic* implementation if one assumes a strict phasic processing of the two representations. Then, instead of having two binary NProcs for 2 *×* 2 distinct values, one such binary NProc would suffice coding during the first phase for the base evidence and then during the second phase for its binding state. Yet, to do so it would require additionally a memory of 1 *bit* to store the information about its base evidence state from the first phase.

**Figure 7:**
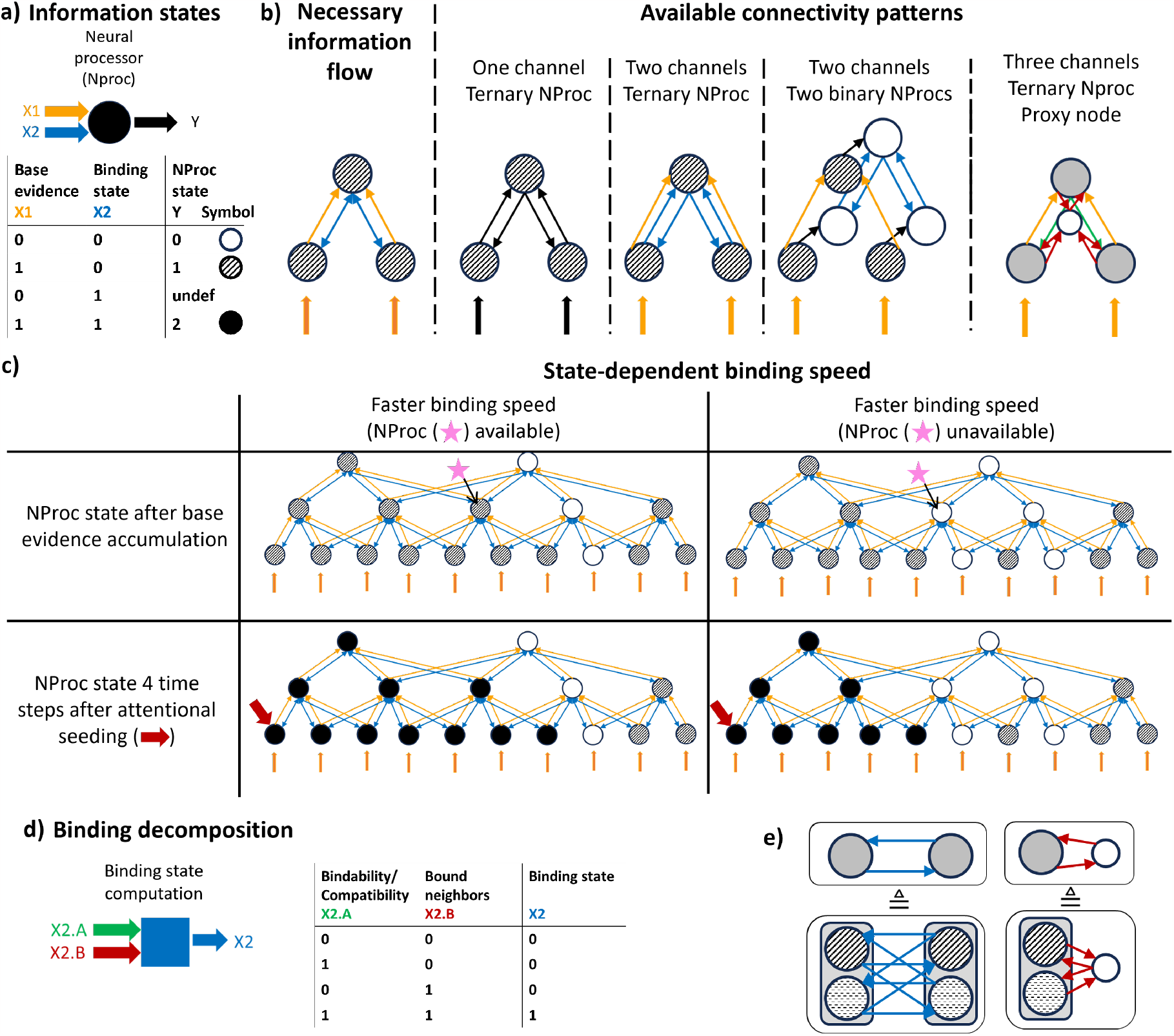
Incremental binding framework and complexity considerations. **a**) Each neural processor (NProc) encodes semantic information states: no evidence (white filling), existing base evidence state (textured filling), or a binding state (black filling). Each NProc captures the processing of a local neural processing site. **b)** Necessary *bi-directional* information flow among NProcs to realize incremental binding (left) and available connectivity patterns to implement this information flow using *uni-directional* couplings (right). The proposed target architecture uses the “Three channels, ternary NProc, Proxy node” pattern. **c)** A hierarchical incremental binding architecture as proposed by [8], for a case of faster (left column) and slower (right column) binding speed, respectively. Here, the case of ternary NProcs is shown. After an initial phase of evidence accumulation (first row) incremental binding can take place among the hierarchy starting from an NProc for which an attentional seed (red arrow) has been provided. Combining base evidence and neighborhood information, specific NProcs among the hierarchy become (un-)available (indicated by pink star) and speed up or slow down the binding process. **d)** Two decomposed sub-signals can be joined by an “and” operation to retrieve the binding state information of an NProc. **e)** Main connectivity variants underlying the complexity considerations. If NProcs are extended for feature selectivities per location (denoted by differet fillings), full connectivity is required to compute the binding state information (left). Using a proxy node to aggregate the binding state information across features greatly reduces the amount of connections (right; see text for details).

**Figure 8:**
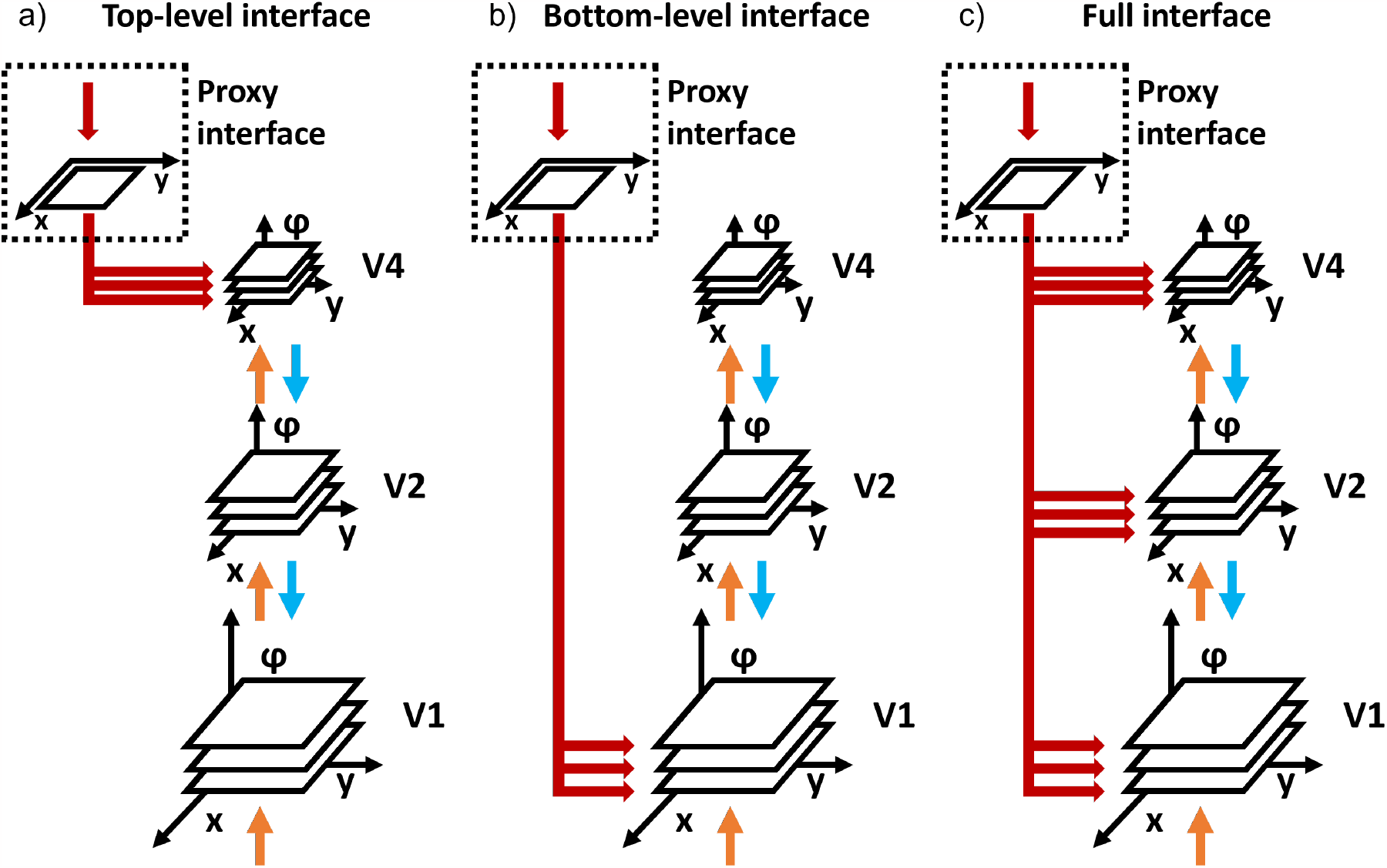
Variants of interfacing with the incremental binding network architecture to proivde an attentional seed from a task module to different hierarchical levels. While a top-level interface (**a)**) restricts provision of the seed of attention to the topmost layer of the hierarchy, a bottom-level interface (**b)**) restricts it to the lowest layer of the hierarchy. Both interface variants form extreme cases of a range of interface patterns. The most general, but also connection-wise most expensive variant is formed by the full interface variant **c)**. Irrespective of the interface variant the number of connections that directly need to originate from and be controlled by a task module depend on the number of features per retinotopic position in the hierarchy. A proxy interface can be utilized as a solution. There, only positional information about the attentional seed is provided by the task module to a proxy map of purely spatial encoding nodes and broadcast to all space-feature encoding NProcs of the associated hierarchical level. The proposed target architecture utilizes the proxy interface pattern in combination with the full interface variant. Additionally, it realizes a bi-directional interaction with the proxy map to realize an information flow pattern which reduces the connectivity complexity for incremental binding further (see text for details). Colors as in Fig. 2.

As a result, the ternary NProc realization is the most efficient one in terms of bits to encode the required representations.

#### 2.3.3. Connectivity schemes and their complexity

To compute their outputs NProcs need access to the signals from other NProcs for base evidence 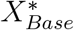 and binding state 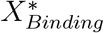 . The transmission of these signals between NProcs happens via unidirectional channels. The necessary connectivity scheme for these channels depends on the signal they transmit. For base evidence signals it is sufficient to connect from NProcs of adjacent scales of the hierarchy (from lower to higher) (Fig. 7 b). As a result, each NProc needs to have channel connections with NProcs from *K*_*l−*1_ neighboring locations from the lower hierarchical scale *l −* 1 and *N*_*F,l−*1_ features per location. For each location, this connection scheme has to be established for all NProcs coding for *N*_*F,l*_ different features. Thus, the required amount of base evidence channels per location is characterized by *N*_*BE,l*_ *∼ O*_*BaseEvidence*_(*K*_*l−*1_ *· N*_*F,l−*1_*· N*_*F,l*_).

For binding state signals the situation is more complex. On the one hand, NProcs need to compute the binding state given the context of their neighbors. This context can be established by channels laterally connecting from neighbors within a hierarchical level, or by feedback connecting from neighbors of the next higher hierarchical level. On the other hand, incremental binding proceeds as a growth-cone-like process. This requires NProcs need to communicate their binding state signals back and forth across the hierarchy to other NProcs that are selective for the same space-feature combination but on another spatial scale (Fig. 7 c):

- If spreading happens faster at a higher hierarchical scale, the corresponding NProcs at the lower scales need to receive feedback to enter the binding state signal as well on their level of representation.
- If spreading happened on a lower hierarchical scale but the local context of an NProc on a higher hierarchical scale would now allow for the binding state, then the lower level’s binding state signal needs to be transmitted to the higher level via feedforward connections.
- If spreading can only continue on the current scale, then lateral connections would be required to signal this information across neighbors and allow them to integrate this evidence with their contextual information.

Thus, the naive realization of binding state channels would require all three kinds of connectivity schemes, feedforward, lateral, and feedback. The required amount of binding state channels per location in layer *l* is then characterized by *N*_*BS,l*_ *∼ O*_*BindingState*_(*N*_*F,l*_ *· P*_*l*_), where *P*_*l*_ = *K*_*l−*1_ *· N*_*F,l−*1_ + *K*_*l*_ *· N*_*F,l*_ + *K*_*l*+1_ *· N*_*F,l*+1_ denotes the pool of *K*_*l*_ neighboring NProcs from the different levels *l* of the hierarchy.

Here, a simplification is possible. It approximates the connectivity scheme by assuming the binding state signal is (de-)composable. Instead of having to compute the binding state signal directly from its neighbors 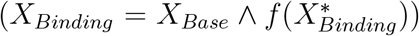, an NProc might compute two intermediate signals for whether it possesses a bound neighbor 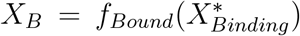 and whether its own space-feature combination is compatible to its local neighborhood neighborhood 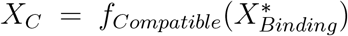. If both signals are available, then the binding state signal can be composed. In boolean logic this would correspond to an “and” function once again 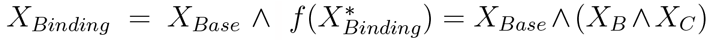 (Fig. 7 d). This simplification assumes independence of the binding state signal and compatibility signal, i.e., the neighbor which provides the information that a binding signal is available doesn’t need to belong to the same neighborhood from which the compatibility of an NProc to the bound representation is calculated.

To extract the information whether a bound representation exists in the neighborhood, the binding state information can be provided by any neighbor irrespective of the feature selectivity. Since it is this binding information which needs to be propagated across all hierarchical scales to establish a growth-cone-like algorithm the decomposition approach can save connections of all three kinds across the hierarchy, i.e., feedforward, lateral, and feedback. What is required, though, is the introduction of an intermediate proxy node, which aggregates the binding state information across features of a local neighborhood to obtain the featureless binding state signal (Fig. 7 b, e). Then, the connectivity complexity can be reduced greatly to *N*_*B,l*_ *∼ O*_*BindingState*_(*N*_*F,l*_ + *P*_*l*_) from *N*_*F,l*_ *·P*_*l*_ (cf. above). Conversely, whether the neighboring bound representation is compatible with an NProc still needs to rely on an evaluation of feature-based compatibility. Yet, by the simplification, it suffices to compute the compatibility information from one of the neighborhoods, i.e. from the next lower, same, or next higher scale, respectively. Thus, the complexity of evaluating the compatibility still needs to be based on evaluating an NProc’s feature against all other features of the neighboring locations, but now the neighborhood only needs to stem from one neighborhood source, e.g. via feedback channels, leading to *N*_*C,l*_ *∼ O*_*Context*_(*N*_*F,l*_ *·* (*K*_*l*+1_ *· N*_*F,l*+1_)) with *K*_*l*+1_*· N*_*F,l*+1_ *< P*_*l*_.

This complexity reduction by decomposition yields the greater benefits in terms of connectivity the more neighbors *{K*_*l−*1_, *K*_*l*_, *K*_*l*+1_*}* each NProc possesses and the higher-dimensional the feature space *{N*_*F,l−*1_, *N*_*F,l*_, *N*_*F,l*+1_*}* is. Thus, these benefits are one of the major motivations for the model architecture proposed in here.

#### 2.3.4. Interfacing demands for incremental binding as a visual routine

An additional requirement for models of incremental binding is imposed by the fact that incremental binding, as a visual routine, needs to be employed within more complex cognitive programs. Such programs are supposed to achieve a behavioral goal, e.g., making a saccadic eye movement to signify a decision about the task. This requires the incremental binding implementation to interface with other routines. Consequently, the question becomes relevant about how efficient the incremental routine can be interfaced. In other words, this question addresses the signature of the routine with its input and output interfaces. Regarding the input, the routine can be triggered by providing it with an attentional seed location. Regarding the output, the routine’s result can be read out in terms of the bound representation.

Under the growth-cone hypothesis, the relevant NProcs to interact with, as targets for input and sources for output, can be found replicated across the different hierarchical scales of the model. Thus, different options exist to realize such interface. Besides the necessity to function under all circumstances, such option should be favorable that requires the least amount of connections. The extreme cases of realization are top-level interface, bottom-level interface, and full interface. The *top-level interface* uses the representation at the highest hierarchical scale of the model as an interface to provide the attentional seed and perform a read-out of the binding state. Since the highest scale offers the most compact spatial representation due to the spatial down-sampling, this realization would yield a low number of connections. The *bottom-level interface* uses the opposing end of the model’s hierarchy as an interface sending information to and reading from the lowest hierarchical scale. Since this is the most extensive spatial representation due to its high resolution, this realization would yield a high number of connections. The *full interface* realization uses all scales of the model hierarchy as representations to interface with. As a result, this realization will yield the highest number of connections. Considering only the number of connections the top-level interface would provide the favorable solution. Yet, there’s a downside: If models should implement the growth-cone hypothesis, then processing of the binding state information can only proceed up to that level of the hierarchy which provides an interference-free representation between the target object’s components and distractors. Likewise, providing an attentional seed via the interface would fail in such cases for which interference would occur at that level to which the attentional seed has been provided. An analogous argument holds for reading out the binding state from such scales. Thus, only a bottom-level or full interface are options for a robust realization of incremental binding under all circumstances.

An additional complexity reduction step that can be applied to either realization consists in the usage of a *proxy interface*. While a cognitive controller would have more degrees of freedom if it would interface with the NProcs of a respective scale directly, it would also need as many direct connections as there are NProcs per feature and location. Yet, if the cognitive control mechanism only needs to interact with spatial information, i.e., to which location the seed of attention needs to be allocated, and to read out which locations are occupied by the bound object, then the cognitive controller doesn’t need such fine-grained control over features. Instead, a proxy interface can encode all spatial locations within a map without a replication across features. It can then provide an interface for the control mechanism and broadcast the attentional seed to all NProcs of a scale coding for the different features at the corresponding location. Likewise such proxy interface can aggregate the binding state from all NProcs of a scale by marginalizing over the feature dimension and only retain the location of the binding states. While such proxy interface comes at the cost of adding another set of nodes for the spatial map representation, it reduces the representational complexity of the interface with which a cognitive control mechanism or other areas of the brain would have to interact.

Consequently, the proxy interface uses *N*_*Inj*_ = *L*_*p*_ + *M* connections to connect the cognitive control mechanism with the layers of the hierarchy targeted by the respective injection method. There *M* = Σ*L*_*l*_ *N*_*F,l*_ is the number of connections needed to connect the interface to NProcs of each space-feature combination across all targeted layers *l* along the hierarchy, i.e., *L*_*l*_ is the number of positions and *N*_*F,l*_ the number of features in a layer. This number *M* is identical to the amount of connections a respective injection method would require without the proxy interface. Conversely, the proxy method needs additional *L*_*p*_ connections to connect the cognitive control mechanism with the interface, where *L*_*p*_ is the number of spatial positions encoded by the interface. Thus, while the total number of connections and nodes increase by *L*_*p*_, the amount of connections which are required from the cognitive control mechanism itself reduce from *M* to *L*_*p*_. This reduction becomes greater the more layers *l* are targeted by the injection method. While the greatest savings are established for the full injection case, already the case of targeting a single layer results in a reduction from *L*_*l*_ *· N*_*Fl*_ to *L*_*p*_.

#### 2.3.5. Resulting model complexity

The proposed model combines the optimal complexity cases for each of the components discussed above and further integrates them with each other. Thus, it establishes normative account for the considerations above. It combines the different optimal components from the considerations of NProc coding, channel connectivity and interface realization. Thus, the presented model uses NProcs with a ternary code mapping the logical states, *noevidence, evidence, bound*, to the gradual output regimes of the pyramidal cell model of zero firing rate, low to intermediate firing rates and high firing rates, respectively. To stabilize these gradual output regimes pyramidal cell model neurons compete via inhibitory pooling across a local space-feature neighborhood. The pyramidal cell model utilizes feedforward channels for base evidence signals entering the pyramidal cell’s basal compartment and feedback channels for the compatibility computation entering the cell’s apical compartment (cf. Fig. 2). The binding state information is stemming from proxy nodes marginalizing binding state information across features per location. It enters into the pyramidal cell computation via a gating component effectively realizing the required “and”-logic. The interface is likewise realized by making use of such proxy nodes. Yet, instead of utilizing a different set of nodes for the interface, a further simplification is performed by using the same proxy nodes for interfacing and model-internal binding state transfrom *L*_*p*_ + Σ*L*_*l*_ to *L*_*p*_. Likewise, the required amount of connections is reduced from *N*_*Inj*_ + Σ*L*_*l*_ *· N*_*B,l*_ *∼ O*(*L*_*p*_ + *M* + Σ*L*_*l*_ *·* (*N*_*F,l*_ + *P*_*l*_)) to *O*(*L*_*p*_ + Σ*L*_*l*_ *· P* ^*\**^), where *L*_*l*_ *· N*_*B,l*_ denotes the fact that the binding state complexity per location *N*_*B,l*_ needs to be considered for the total amount locations *L*_*l*_ per layer *l* and the multiplication mission. Therefore, the required amount of proxy nodes is further reduced by *L*_*l*_ by *P* ^*\**^ = (*N*_*F,l*_ *· K*_*p,l*_ + *N*_*F,l*_ *·K*_*l,p*_) *< P* denotes the fact that the different layers *l* now need to connect to the common proxy map *p* via a local neighborhood that maps their spatial scale and vice versa, by *K*_*p,l*_ and *K*_*l,p*_, respectively, instead of mapping to each adjacent layer. This results in a single proxy map, located in the interfacing module, that serves to encode binding state information and to signal it to NProcs of corresponding positions across the hierarchy and to other brain areas involved in functions such as cognitive control or occulumotor behavior.

## 3 Discussion

Here, we described a mechanistic model of incremental binding that is able to successfully explain a broad range of experimental evidence and implements the growth-cone hypothesis. The model representations during simulation matched with neural correlates reported in the literature, replicating phenomena of constant up-modulation to signal a task-relevant object, distance-dependent onset of up-modulation, and growth-cone-like binding speed variations in presence of distractors. Furthermore, lesion studies identified the causal involvement of specific connection types. Direct feedback connections between larger scales of the hierarchical model are necessary to keep the growth-cone-like incremental binding properties intact. Without those connections binding was still able to proceed, but at an almost constant, lower speed. Beyond these findings, the model is also able to handle intersection cases between a target and a distractor curve extracting grouped objects that align with the Gestalt law of good continuation.

To achieve these results, the model only needs a single type of neural processors across the hierarchical layers of the visual cortex module and utilizes a common representation for providing output to other nodes in the network. Neurons of the interfacing module filter the responses of neurons from the visual cortex module for their binding state by accumulating their output and thresholding it against a high enough value. This threshold mechanism is filtering the neural responses effectively for task-relevance. Ideally, only if the neurons reside in a regime of up-modulated activity, their values will be high enough, so that their activity will cross the threshold and will be permitted into the output processing of the location within the interfacing module. A task module serves as a cognitive control mechanism and provides external input to the interfacing module. This external input takes the form of an attentional seed location and is triggering the incremental binding process between the visual cortex module and the interfacing module.

These design decisions for the model can be contextualized within a theoretical framework of incremental binding. Within this framework we identified constraints and evaluated the individual model components against possible alternatives regarding their connectivity complexity, coding efficiency, and implemented logic. Finally, this complexity analysis of the model and its components showed that the model compares favorably against other possible alternatives, which could have likewise been used as structuring principles of such an incremental binding architecture. Thus, it makes the proposed model a good candidate for a normative account of incremental binding.

### 3.1. Incremental Binding could be explained by an interplay of cortex and thalamus

Based on the proposed detailed computational model architecture, the question arises how the computational structures may relate to the visual cortical anatomy. The spatial hierarchy of representational layers best corresponds to visual areas along the ventral pathway, as commonly suggested [38, 39, 40].

Neural processors within each layer model pyramidal cells and their local inhibitory circuits. The model neurons provide a close functional description of the underlying biological neuron’s computation. They capture main findings, namely, asymmetric combination of feedforward and feedback input via apical-basal integration [41], thalamic gating of respective integration [42, 43], and response up-modulation through attention [9]. That pyramidal cells play a dominant role for cortical computation is likewise suggested throughout experimental and computational approaches (see [44, 45] for an overview). Since the model’s framework provides a clear guidance for the distinct roles of each connection type, one yields interpretable neural representations. A direct mapping can be made between the concept of a binding-state-dependent compatibility computation and the apical compartment of the model’s neural processor. Thus, the model provides a prediction in assigning such role to the apical dendritic integration of pyramidal cells.

Another important component of the model consists in the interfacing module that implements a neural sheet of proxy nodes, serves as an interface between hierarchical layers of the visual cortex module, and gates the apical-basal integration of pyramidal model cells. We suggest that this proxy node computation is best matched by higherorder proportions of the visual thalamus, such as the pulvinar. First, given recent neurophysiological evidence [43], gating apical-basal integration of pyramidal cells is best explained by thalamic input sources. Second, thalamus provides a good fit to the interfacing role, as thalamus and cortical areas are reciprocally connected in a shallow hierarchy [34]. Third, in line with our lesion studies, the relationship between thalamic and cortical activity is a causal one [46]. And fourth, the featureless, task-dependent representation employed by the proxy nodes of the interfacing module, fits well with the identified role of thalamic function in attentional selection [47, 48]. While the evidence about the thalamic representation might not be completely clear yet regarding whether its rather featureless or coding for specific features, like visual motion or motor information [49, 50], overall the representation seems to be involved in task-related processing and lower-dimensional than that of the associated visual cortical areas, which is are crucial factors of our interfacing module.

Regarding the thalamic influence on the apical-basal integration process we modeled it by a direct coupling onto the pyramidal cell. Yet, it might likewise be possible that the thalamic gating role is rather mediated indirectly via apical (dis-)inhibitory circuits [51, 52, 53], which would describe an extension to the proposed model rather than a contradiction.

In light of evidence from the past decades, the role that thalamus is thought to play in cognition has changed from being a mere relay station that duplicates (sub-)cortical signals to a more important one that is involved in coordination [54, 55]. Yet it’s precise functional role is still debated and led to a range of (non-)exclusive suggestions [56]. Suggestions closest to the original relay station idea assign the thalamus a function of selectively (de-)activating signals that are passing through this trans-thalamic route. Such selective choice of signal propagation would make the thalamus a *gatekeeper* [57, 58], *or switchboard* [59]. Similar to this is the idea of the thalamus as a *hub* [60, 61, 62]. There, it serves to integrate different signal streams and promotes these further to other brain regions and potentially plays a role in multiple cognitive functions. Focusing further on the integrating behavior of the thalamus, but for more specific cortical functions, it has as well been suggested that thalamus serves as a blackboard [63, 64]. Under such blackboard assumption cortical processes can write information to the thalamus in order to store and update information and read from it in order to communicate with other processes. In this regard, the thalamic role would be in storing current system state information and providing a buffer that exposes a common interface to different cortical areas which take part in executing a common function. A similar blackboard role has as well been suggested for primary visual cortex [65, 66]. Thus, it could be likewise possible that the blackboard function is established in concert by visual cortical and thalamic counterparts. In a broader view of integrating processes across cortical sites it is also suggested that thalamus serves as a gateway, or hub, for permitting information and regions into a conscious state [67, 68, 69, 70]. Focusing rather on behavioral implications, the thalamus could play as well a *task-related role*. E.g., pulvinar function was found to serve as a confidence map which codes for the confidence in an observer’s decision [47].

In our model the thalamic regions are modeled by the interfacing module and are assigned properties of task-related coding, interfacing, and blackboard functionality. In terms of a *task-related role*, it aggregates coarse-grained location-specific information about task-relevant integrated representations from cortical pyramidal cells. In terms of an *interface*, it can be used as such for attentional control, as the task module can provide a seed location of attention to the visual cortex module via the interfacing module’s proxy nodes. And in terms of a *blackboard*, the interfacing module contains the binding state of the visual cortex module, aggregates it in a map, only provides supra-threshold activity for those positions that signify binding states. Thus, it takes part actively in the binding process and exerts cognitive control onto the cortical architecture based on task-related guidance signals.

Where these task-related signals arise from before being passed to the thalamus, and whether those places themselves represent the locus of attention in a similar spatial map, is outside the scope of the model. Many structures of the brain have been reported to be involved in attention, but in how far each plays a causal role is not clear yet [71, 72]. Yet, reasonable cortical candidates exist that could represent such locus of attention and relay it to visual cortex through thalamus. These candidates can either be found in prefrontal regions, such as the frontal eye fields (FEF) [73, 74, 10], posterior regions, such as intraparietal sulcus (IPS) [75], or also in temporal areas [76], such as the dorsal part of the posterior inferotemporal cortex (PITd) [77] . Additionally, that other sub-cortical sites, such as the SC (Superior Colliculus), further modulate this processing by thalamic projections is likewise feasible [78, 79].

While the above evidence suggests that especially higher-order thalamic sites play a relevant role in cortical attentive function, it nevertheless cannot be ruled out, that the role of the model’s interfacing module isn’t associated to higher-order thalamic sites, but to other cortical areas directly. Yet, in this case the specific connectivity pattern as implied by the model is rather unlikely. There, the cortical area in question would need to connect to each apical dendrite of the visual areas’ pyramidal cells and subserve a specific gating role. This gating signal would need to be distinct from the contextual evidence aggregation role that other apical feedback inputs fulfill, and therefore, would likely need a different kind of connection type.

Thus, given our findings, we suggest that incremental binding might best be explained by an interplay between cortex and higher-order thalamus.

### 3.2. Broader implications of the complexity considerations

The investigations about the complexity of different computational motifs for incremental binding and their connectivity patterns can serve a broader purpose beyond judging the proposed model’s complexity. Our additional assumptions and constraints extend an earlier framework for algorithmic descriptions of incremental binding [25, 8]. The framework allows to reason about incremental binding on an algorithmic level and our complexity considerations can help identify open questions to guide modeling decisions. Furthermore, existing models can be analyzed with respect to those decisions and how well they fit a normative account regarding their coding, representaitonal and connectivity complexity. Dependent on the model these decisions are either made explicitly or implicitly. When it comes to coding for the different states of a node, different choices are possible. These choices can be, e.g., introduction of a separate population and/or channels for the incremental representation versus the base representation [17, 80, 21, 18, 19, 22]. Likewise, a multi-step approach could be a solution to provide distinct phases of base evidence accumulation vs. incremental binding propagation [6, 23]. Furthermore existing learning-based architectures, for which the relationship between different pathways and neurons in the network and their representational role might be less clear, could be analyzed in this regard [81, 82, 83, 24].

Unlike the previously mentioned models, the currently proposed model doesn’t foresee completely separate populations for the base representation and incremental binding state, nor does it require phasic processing. Instead, it utilizes two-compartment neural processors and a by-pass route via the interfacing module, that specifically filters for binding state information.

Investigating all the different models further with the tools used here, e.g. w.r.t. computational and representational complexity, could provide further insights into the conceptual strengths and weaknesses of the respective models. Likewise, these comparisons could provide further insights into which model choices lend themselves for easy trainability within deep learning frameworks and shed light on why that might be the case. Furthermore, by providing numbers in terms of representational, as well as computational complexity, models also become more comparable. There, it might become more apparent how big of an implementational overhead exists per model in comparison to an ideal realization.

Despite the already discussed cases for how to encode the different base and binding states, further options in the design space exist. These options offer modelings solutions, which have been unexplored so far. For example, the same neural processor could code for gradual base evidence via tonic firing rates and switch into a propagating binding state information by providing higher-rate bursting output. There, different recipient nodes could filter the neuron’s output for lowvs. high-frequency codes to provide multiplexing among a single neuron’s output channel [84].

### 3.3. The broader context of binding and visual-cognitive architectures

The investigated architecture and proposed framework focused on incremental binding mechanisms in visual grouping. Nevertheless, one of the underlying assumption for all the considerations has been, that visual cortex is capable of flexible processing dependent on task demands and attention. Thus, it shouldn’t only be able to perform incremental binding, but can engage in different tasks, such as grouping, searching, comparing, or tracking objects.

This generality and flexibility has been discussed in the context of visual routines [4, 5] and cognitive programs [6]. The common idea is that, similar to a computer, visual cortex can make use of different encapsulated computational motifs to find solutions to specific tasks, and that more complex tasks can be tackled by compositions of these simpler motifs. The model proposed in here implements one such motif, namely incremental binding. On a level of elemental operations, it therefore needs to be able to read data, e.g. from the task-module, store data in the interfacing module, and perform a recursive binding operation starting with the attentional seed and ending when no further increment to the grouped object is encountered. What is missing so far in terms of visual routines and cognitive programs for the proposed model is the flexible combination with other routines and a task-dependent selection of the respective routines. Nevertheless, the proposed model provides a clear interface for possible interaction with a cognitive control mechanism and for exchange of intermediate with results with other routines, and doesn’t possess any hard-coded components that would restrict its function purely to incremental binding.

To achieve the computational operations for incremental binding the system is evolving in a decentralized and parallel fashion. Yet, the incremental binding process unfolds only sequentially in time and localized in space. It depends on the availability of local neighbors for compatibility computations and on a gating process that is likewise dependent on a local neighborhood. This parallel-yet-sequential processing has similarities to earlier such ideas [85, 2, 86, 3], where decentralized processes operate in parallel until a sequential bottleneck forms a decision, that then can lead to the next step of a parallel execution process. For the case described in here, this formation of a rather discrete decision (bound vs. not bound) is happening by the accumulation process within the interfacing module. It accumulates the evidence for a binding state at the respective position across a decentralized pool of neurons in the visual cortex module. Dependent on the state of this accumulation the interfacing module then gates the parallel and decentralized apical-basal integration processes of the decentralized pool of neurons among the hierarchy of the visual cortex module.

On a broader scope, the model is compatible with theories of cognitive architectures. E.g., the model’s interfacing module could be interpreted as a buffer of visual information in terms of ACT-R [87]. Likewise, the flow of task information in the model, where a task-module triggers a computational motif in the visual cortex module and would later on be able to read out the resulting attended object, fits the view of a parietal-prefrontal processing paradigm [88]. Based on the employed apical-basal integration mechanism that is gated by higher-order thalamic input [43] the model of pyramidal cell computation proposed in here fits theories about visual cortex and thalamo-cortical interaction in consciousness frameworks [89, 70].

Relating the model to other theories of binding and object-based attention commonalities and differences can be found. In comparison to the feature integration theory (FIT) [27], the model wouldn’t represent specific stimulus features, such as contour or color in a decomposed fashion, rather they would be assigned to a common high-dimensional feature space in visual cortex module, where different neurons would have different feature selectivities. Since, so far the model experiments were only concerned with processing oriented contrast information, i.e. line segments, this distinction from a decomposed representation of feature spaces would only become clear in more sophisticated model versions. A component in common with FIT is the existence of a spatial relevance map that helps guide the selection of what places should be bound. In the proposed model this role is assigned to the task module, which mediates this spatial relevance via the interfacing module to establish the incremental binding in the visual cortex module. In comparison to a more recent proposal for an architecture of attention [33], the model’s interfacing module possesses some commonalities with the idea of gain maps. Gain maps represent a spatial projection from the hierarchical stream of layers that perform feedforward-feedback interactions and back-project this representation to selectively influence the respective hierarchical processing in favor of the map’s content. This motif of selectively biasing processing is also found in the model’s interfacing module. It does so by representing a spatial projection of binding state information from the visual cortex module and back-projecting this information onto a local neighborhood across the hierarchy. Yet, the interfacing module mostly is concerned with mediating this process of selection and could likewise be influenced from other gain-map-like areas that influence the selection process further (see the above discussion about candidate regions for signals of attentional control).

### 3.4. Limitations and Possible Extensions

The framework and model so far have been developed to investigate incremental binding processes operating on oriented contour segments. Beyond such evidence, a broader set of reported experiments for surface-related and object-based binding exists [90, 91, 11]. Thus, an interesting future direction consists in extending the framework with respect to more complex feature spaces, such as surface representations [92], or object(-part)-level representations [7], which might even be encoded in non-retinotopic spaces. A specific example of a more complex interplay between features consists in a version of the curve tracing task which is based on motion coherence stimuli [12]. It likewise potentially requires spreading of attention. Yet, while a model for this task exists [93], it remains unspecific with respect to the incremental versus parallel nature of binding and an interplay of these mechanisms and invites for further investigations using the model proposed in here.

Furthermore, the proposed model has been designed with keeping in mind the necessity of providing an efficient interface for incremental binding giving rise to a normative account regarding coding, connection and representation complexity. Thus, it opens up an venue for investigating the model’s capabilities in regard of other visual-cognitive processing, i.e., visual routines. One such example would consist in using the architecture and the theoretical complexity analysis to investigate visual search, which is another candidate often used for reasoning about binding in the brain [94, 95, 96]. A resulting model might then be able to implement both, visual routines for incremental binding and visual search.

Regarding the modeling of thalamic function in thalamo-corticcal interactions, it would be of further interest to investigate relations to other existing relevant models. Logiaco et al. show how exerting control over cortical computational motifs can be efficiently realized via a smaller population of thalamic neurons [97]. There neurons are modeled in less detail where rather their role within a pool and the connection density are the guiding principles. Munn et al. show how the formation of state-dependent pyramidal cell bursting can be linked to thalamo-cortical interaction [98]. They provide an information-theoretic investigation and link the thalamo-cortical interaction to a maximization of integrated information in pyramidal cell computation. Remaining on a finer physiological level, the simulations, yet, don’t imply a specific algorithmic function, but rather provide an argument for optimized information transmission. In a related large-scale model for more abstractly investigating attractor dynamics of thalamo-cortical interactions, Müller et al. show how thalamus can play an important role shaping the cortical information state regarding consciousness [99]. Investigating interrelations between these models and the work presented in here could contribute to further the understanding of thalamo-cortical interactions and the modeling thereof.

Regarding our complexity considerations, an additional topic of interest, which might guide future models for (incremental) binding, is learning. Considering how easy or hard it is to learn, or evolve, specific connectivity structures and parameter combinations could help judge the biological plausibility of different models. On a similar note, it might be useful to further investigate the power of potential binding mechanism of our model in the context of deep learning. For example, within the scope of contextual binding [100] or segmentation challenges [81, 23]. There, it also could further be compared to other learned models of incremental binding [21, 81, 23, 22].

Another interesting route for future investigation could consist in further challenging the complexity analysis and/or existing models concerning, e.g., the growth cone hypothesis. For example, the analyzed attentional seed injection methods investigated in Sect. 2.3.4 identified a shortcoming for direct top-injection. Interference cases between the target object’s representation and a nearby distractor renders this case ineffective prohibiting backprojection of the binding signal. Yet, maybe a solution could exist if, e.g., the investigated feature space becomes high-dimensional enough, such that the interference is resolved simply by the sparsity of the high-dimensional embedding. Testing such ideas also experimentally might shed further light on how the brain implements incremental binding on a neural level.

## 4. Materials and Methods

### 4.1. Model details

#### 4.1.1. Network model

On a network-level, the model can be viewed as a graph of nodes (neural processors) and channels. The nodes are arranged in retinotopic space and code for certain input features, i.e. orientations, forming feature maps of a model area. The areas of the visual cortex module form a scale space and have a relative size with respect to the input of [1, 0.5, 0.25] for the layers V1, V2, and V4, respectively. The interfacing module likewise forms a retinotopic map, but only with a single feature channel, i.e., binding state. The interfacing module’s relative size with respect to the input was set to 0.66 thus ranging between the spatial resolution of V1 and V2. The channels connecting the neural processors are modeled by localized interaction kernels. The kernels abstractly represent underlying neural connectivity patterns and provide weighted connections of node activities from a local spatial neighborhood around the target node’s position. While being localized in the retinotopic space, kernels provide connections to all feature nodes within this localized vicinity. Updates among these connections are propagated by computing the convolution across space of the respective kernel with the source nodes. Overall, the application of a kernel is performed by

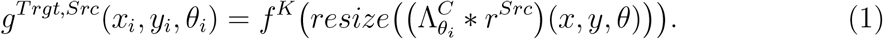

Each node has a certain retinotopic position *x*_*i*_, *y*_*i*_ and a feature value *θ*_*i*_ it is coding for. Here, 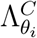 describes the kernel of channel type *C* that represents channels connecting the source node *Src* with the target node *Trgt* based on the source node’s activity level *r*^*Src*^. The operator denotes a convolution with circular boundary treatment among the feature dimension and zero-padding among the spatial dimensions. To adjust for the different spatial sizes of the different layers a resizing operation *resize* is applied after each convolution using linear interpolation in space. To restrict the range of input values to each node, the convolution operation is followed by a non-linearity *f* ^*K*^ = *min*(*max*(*x*, 0), 1).

The overall connectivity scheme of the network is given by Table 1. Within the network, four types of channels exist: feedforward (FF), feedback (FB), and proxy-related, with a distinction into channels projecting to (B1) and channels projecting from (B2) the thalamic interfacing module. The local pyramidal cell processing of a neighborhood is aided by an inhibitory (Inh) interaction.

**Table 1:** Network connectivity structure.

| From \ To | Inp | V1 | V2 | V4 | HO | S |
| --- | --- | --- | --- | --- | --- | --- |
| Inp | - | FF | - | - | - | - |
| V1 | - | Inh | FF | - | B1 | - |
| V2 | - | FB | Inh | FF | B1 | - |
| V4 | - | - | FB | Inh | - | - |
| HO | - | B2 | B2 | B2 | - | - |
| S | - | - | - | - | B1 | - |

The respective kernels between nodes of the hierarchical layers are space-feature separable and defined by

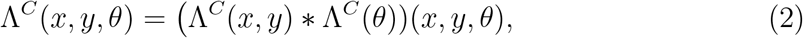

where Λ^*C*^(*x, y*) defines the kernel component in retinotopic space and Λ^*C*^(*θ*) the kernel along the feature dimension. There, kernels of the feedforward channel *C* = *FF* consist of a Gabor kernel for the spatial component

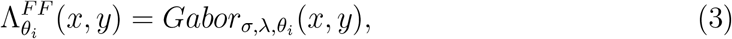

with a standard deviation *σ* for its envelope, wavelength *λ* = 2*\*π*σ*, and orientation *θ*_*i*_, and a Dirac kernel along the feature component

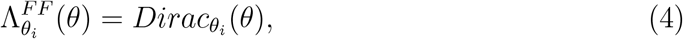

for a given target node’s orientation *θ*_*i*_.

Kernels of inhibitory type *C* = *Inh* consist of a Gaussian kernel along the spatial dimension

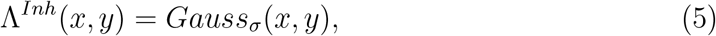

with standard deviation *σ* and a uniform kernel along the feature dimension spanning all feature channels, i.e. size *ρ*_*θ*_ = *n*_*θ*_

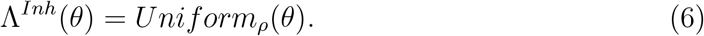

The composite inhibitory kernels have been normalized to a sum of 1.

Kernel of feedback type *C* = *FB* consist of a Gabor kernel along the spatial dimension (Eq. 3) and two feature kernels that are separately convolved with the positive and negative values of the Gabor filter respectively (Eq. 2). While the feature kernel for positive values is given by a rectified cosine half-wave

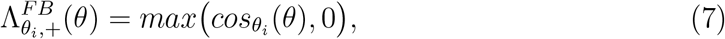

the feature kernel for negative values 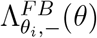 is a uniform kernel (Eq. 6). The feedback conductance is computed by filtering the input from the next higher layer with each of the kernels 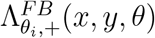 and 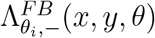 separately and afterwards adding up the result:

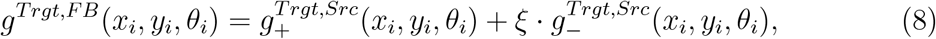

where *ξ* is a scaling factor for the negative contribution of the feedback signal. Gabor filters for both, *FF* and *FB* connections, had a phase shift of 0 (even component), an aspect ratio of 1, and had their positive and negative values normalized to sums of 1 to compensate for the DC component. To avoid cut-off artifacts matrix sizes of the Gaussian and Gabor kernels were set to 1 + 2 *·*5*σ*.

The kernels connecting the hierarchical layers of the visual cortex module with the interfacing module are defined as follows. Binding channel connections of type *C* = *B*1 connecting a hierarchical layer source *Src ∈ {V* 1, *V* 2, *V* 4*}* to the interfacing module *Trgt* = *HO* are given by a composite kernel (Eq. 2), where the spatial component, as well as the feature component are defined by uniform kernels (Eq. 6; with limited extent in space *ρ*_*x*_ *× ρ*_*x*_ and spanning all feature channels *ρ*_*θ*_ = *n*_*θ*_). The binding channel connections of type *C* = *B*2 connecting the interfacing module *Src* = *HO* to a hierarchical layer target are purely spatial kernels, as the interfacing module doesn’t code for any feature *θ*, and are likewise defined by uniform kernels (normalized to sum of 0.5) with limited spatial extent (Eq. 6).

#### 4.1.2. Pyramidal cell model

Within the network each node represents a neural processor that abstractly captures the computation of a local neural microcircuit of finer detail. For the hierarchical layers’ nodes, this microcircuit can represented by a cortical column model [101]. We describe its computation by means of a pyramidal cell, which is thought the be most relevant processing element of such microcircuits [44, 45]. The pyramidal cell model consists of two dynamical equations describing the development of basal *b* and apical *a* membrane potentials:

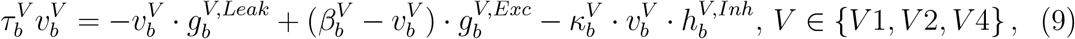

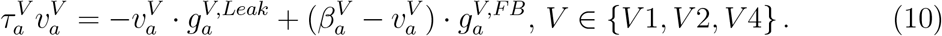

Such cells reside on all layers *V* of the model hierarchy, i.e. areas V1, V2 and V4. These conductance-type equations capture the integration of basal, peri-somatic influences on the cell’s membrane potential and the changes on apical distal dendritic potentials. Here, constants *β*^*V*^ describe the excitatory reversal potentials, which provide an upper bound to the membrane potential values, *κ*^*V*^ is a scaling constant for the divisive inhibitory influence, and time constants *τ* ^*V*^ steer the time scale on which the evolution on which the temporal dynamics take place. Conductances *g*^*V,T*^ describe input-dependent influences on the potential based on specific input channel types *C FF, Inh, FB, B*2 . The inhibitory component *g*^*V,Inh*^ is mediated by a nonlinearity *f* ^*Inh*^ = 1.3*/*(1 + exp ((*x* 0.25) 20)), which captures the response behavior of local inhibitory interneurons

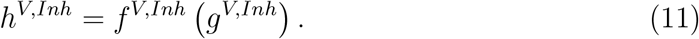

Additionally, *g*^*V,Leak*^ describes a constant leakage, and the excitatorily driving conductance *g*^*V,Exc*^ is defined by

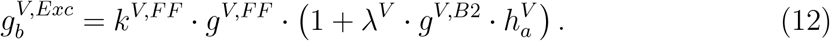

This equation describes the three-way interaction between the feedforward channel, the feedback channel, and the interfacing module input, where *k*^*V,F F*^ and *λ*^*V*^ are scaling constants. The feedback channel’s influence is mediated by the apical component via

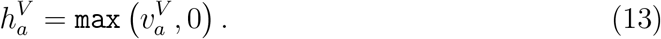

The three-way interaction among these inputs is asymmetrical (cf. Fig. 2). While feedforward input *g*^*V,F F*^ is able to provide on its own an excitatory on the basal membrane potential, feedback input 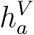 only takes effect if it coincides with interfacing module’s gating input *g*^*V,B*2^. In addition, the feedback’s effect on the excitatory conductance is of a gain enhancement type. So, it will up-modulate existing feedforward input (abstractly, i.e., *FF* + *FF ·FB*; [101]), but will take no effect if no feedforward input is present. For the highest layer of the hierarchy, V4, no feedback exists (cf. Tab. 1). As a result the apical membrane potential will not receive any input, and thus, won’t influence the cell’s basal dynamics.

To compute the output rate *r*^*V*^ of the of the pyramidal cell its basal membrane potential 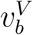 is passed through a non-linearity *f* ^*V*^ = *min*(*max*(*x*, 0), 1)

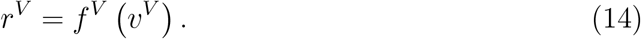

#### 4.1.3. Thalamic cell model

Nodes of the interfacing module describe the processing of higher-order thalamic model neurons. The evolution of their membrane potential *v*^*HO*^ is governed by the differential equation

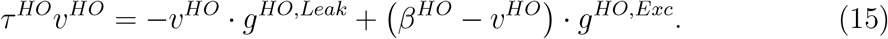

The equation is structurally similar to Eq. 9 and Eq. 10 with a constant leakage conductance *g*^*HO,Leak*^, an excitatory reversal potential *β*^*HO*^, a time constant *τ* ^*HO*^, and an excitatory conductance term *g*^*HO,Exc*^. For thalamic cells, this term consists of a summation across multiple inputs *V* according to

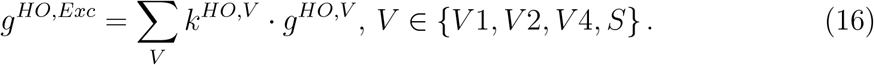

This summation is weighted by constants *k*^*HO,V*^, *V ∈ {V* 1, *V* 2, *V* 4,*} S* and describes the thalamic evidence accumulation of binding signal information coming from the different neural processors of the visual hierarchy via channels of type *C* = *B*1 and other sites of the brain *S* providing a seed of attention. In this case *S* is the task module providing input to the interfacing module. The accumulated evidence (Eq. 16) is mediated by the temporal integration (Eq. 15) and then transformed into the cell’s output rate *r*^*HO*^

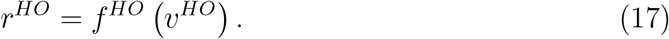

The non-linearity *f* ^*HO*^ = 1*/*(1+exp (*−* (2 *· x−* 1) *·* 50)) is of sigmoidal kind and rather steeply segregates non-binding from binding-states by its threshold and, therefore, provides a filtering mechanism for up-modulated activity coming from the visual cortex module. Since the input aggregation scheme (Eq. 16) is purely linear, some drawbacks can exist depending on the parametrization of the model. I.e., if the up-modulation effect is too small, then the sum of aggregated activities stemming from up-modulated neurons at a site of lower overall base-level activity might be smaller than the sum of aggregated activities stemming from base-level activated neurons at a site of higher overall base-level activity. In this case the interfacing module wouldn’t be able to properly detect the bound locations. On the one hand, this issue didn’t occur for the simulations using the proposed parametrization. On the other hand, this weighted summation could easily be replaced by, e.g., a maximum selection among the inputs to counter the issue. Yet, we omitted such countermeasure for the sake of a more simplistic model which was still able to capture the observed incremental binding correlates.

### 4.2. Details on Experiments

Experiments have been performed using the Julia programming language^1^.

#### 4.2.1. Panel creation

The input data *r*^*Inp*^ for the experiments have been created in two steps. First, a contour of respective shape has been defined on a pixel grid with a line width of 1 pixel and a grayscale intensity value of 1. To have a smoother transition for diagonal elements two neighboring pixels have been set to intensity values of 0.75 instead of setting one pixel in an 8-neighborhood to 1. Second, the resulting image has been padded by 8 pixel to each side, blurred using Gaussian filtering (*σ* = 1.0) and subsequently up-scaled by a factor of 3 to the desired input size followed by linear interpolation. All input images have been normalized to a maximum of 1. For the case of varying distances between target and distractor curve, the wide distance segment had a spacing of 8 empty pixel, and the narrow distance segment had a spacing of 4 pixel. The vertical extent of the lines was 81 pixel. The proportion of narrow vs. wide segment length has been varied parametrically between 0.15 and 0.85, which provided a range in which the stimulus properties stood intact and a constant overall curve length was maintained. The extent of the narrow segment has been positioned symmetrically around the contour’s longitudinal center.

For the crossing stimuli the same procedure has been applied. The spacing between the two lines in the defined pixel grid was 11 pixel and the straight line 42 long.

The attentional seed values have been provided via a separate map *g*^*HO,S*^. There, the creation procedure consisted of pixel-precise definition, no blurring, and up-scaling. Each seed of attention has been provided as a single pixel of intensity value 1 at the corresponding end point of the respective contour.

#### 4.2.2. Simulation details

All simulations have been performed with the same set of model parameters. An overview of the parameters can be found in Table 2.

**Table 2:** Parameters used throughout the different incremental grouping experiments.

| Parameter | Value | Explanation |
| --- | --- | --- |
| **V1-V4** |  |  |
| $n_\theta$ | 12 | Number of orientation feature channels |
| $g^{V,Leak}$ | 1.0 | Leak constant, $V \in \{V1, V2, V4\}$ |
| $\beta^V$ | 1.0 | Excitatory reversal potential, $V \in \{V1, V2, V4\}$ |
| $\kappa^V$ | 7.5 | Divisive inhibition factor, $V \in \{V1, V2, V4\}$ |
| $\lambda^V$ | 15.0 | Apical impact factor, $V \in \{V1, V2\}$ |
| $k^{V1,FF}$ | 0.25 | Re-scaling input activity |
| $k^{V,FF}$ | 3.0 | Re-scaling activity between layers, $V \in \{V2, V4\}$ |
| **HO** |  |  |
| $g^{HO,Leak}$ | 0.7 | Leak constant |
| $\beta^{HO}$ | 1.0 | Excitatory reversal potential |
| $k^{HO,S}$ | 8.0 | Input scaling factor |
| $k^{HO,V1}$ | 16.0 | Input scaling factor |
| $k^{HO,V2}$ | 4.0 | Input scaling factor |
| $k^{HO,V4}$ | 1.0 | Input scaling factor |
| **Kernels** |  |  |
| $\sigma^{V,FF}$ | 2.0 | envelope standard deviation, $V \in \{V1, V2, V4\}$ |
| $\sigma^{V,Inh}$ | 1.0 | kernel standard deviation, $V \in \{V1, V2, V4\}$ |
| $\sigma^{V,FB}$ | 2.0 | envelope standard deviation, $V \in \{V1, V2\}$ |
| $\xi$ | 2.5 | Inhibitory feedback component scaling factor |
| $\rho_x^{HO,V}$ | 3 | Spatial extent of kernel, V to HO, $V \in \{V1, V2, V4\}$ |
| $\rho_x^{V1,HO}$ | 5 | Spatial extent of kernel, HO to V1 |
| $\rho_x^{V2,HO}$ | 9 | Spatial extent of kernel, HO to V2 |
| $\rho_x^{V4,HO}$ | 13 | Spatial extent of kernel, HO to V4 |

The listed parameters were determined by manual search. While the parameter choice proved itself robust for the given experiments, no dedicated robustness tests have been performed. Yet, during optimizing for these experiments, the parameter regime was graceful around the chosen values, so that the choice of integer numbers was often reasonably good enough. Vice versa, this means that a computationally more demanding optimization of the parameters could yield even better quantitative fits to experimental data.

To keep the computational cost for simulating the experiments low, steady state iteration has been performed for all dynamical equations instead of a more detailed numerical integration scheme. For all simulations all state values have been initialized with zeros.

#### 4.2.3. Metrics and Evaluation Schemes

To evaluate the simulation results, the following metrics and computations have been performed.

The modulation index *MI* has been computed by comparing the neuron’s activity between two conditions during each simulation, respectively: the baseline condition at the beginning of the simulation *base*, where the respective neuron still was in a base representation state before the incremental binding started, and the condition at the end of the simulation *att*, when the respective neuron has been labeled by attention.

The neural activities during these conditions,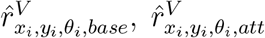, served as firing rate estimate to be used for computing the modulation index. The modulation index has then been computed according to [9]:

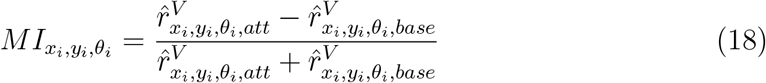

Computing the time until the complete curve has been up-modulated by attention, the modulation onset time of the last neuron position in V1 along the curve’s outline has been used. The modulation onset time was defined as the time step at which the neural activity crossed 50% of its maximum that occurred during the respective simulation. The onset time has been determined for the population’s activity summed along the feature dimension *θ* to obtain a single onset time per retinotopic position. To estimate the relationship between a variation of the proportion of narrow segments along the contour this proportion has been varied and linear regression was performed across the completion times of the different simulation runs.

The computation for the speed profiles in the growth cone investigations has been based on the same modulation onset time definition as described above. Yet, to compensate for pixel discretization artifacts and other numerical issues the onset times have been smoothed across neighboring neuron positions along the contour outline. Smoothing has been performed by convolving the modulation onset times with a binomial kernel Λ_*s*_ = [1, 4, 6, 4, 1] */*16.0. Then speed values 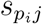 for positions *p* along the contour outline have been computed by taking the pair-wise difference between smoothed onset times of neighboring retinotopic positions *p*_*i*_, *p*_*j*_ and subsequently taking the inverse of this onset time difference, i.e.,

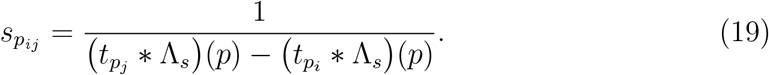

In theory the denominator of this equation can become zero or even to lead negative speed estimates, if the smoothed time estimates for indices *i* and *j* don’t increase monotonically. In practice, though, such monotonicity for the indices along the contour outline has been observed during the simulations.

To compute estimates of the speeds per contour segment (first wide segment, narrow segment, second wide segment) linear regression was performed for the modulation onset times of the neurons per segment. The inverse of the determined slope factor then yielded the speed estimate.

## Acknowledgments

The authors received no specific funding for this work.

## Author Contributions

The authors made the following contributions according to the CRediT taxonomy:

*Conceptualization*: DS, HN

*Data curation*: DS

*Formal Analysis*: DS

*Funding Acquisition*: HN

*Investigation*: DS

*Methodology*: DS, HN

*Project administration*: DS, HN

*Resources*: HN

*Software*: DS

*Validation*: DS

*Supervision*: HN

*Visualization*: DS, HN

*Writing - original draft* : DS

*Writing - review & editing*: DS, HN

## Footnotes

1 Code and data are available online: https://github.com/schmidDan/incremental_binding

